# Dynamin-related Irgm proteins modulate LPS-induced caspase-4 activation and septic shock

**DOI:** 10.1101/2020.03.18.997460

**Authors:** Ryan Finethy, Jacob Dockterman, Miriam Kutsch, Nichole Orench-Rivera, Graham Wallace, Anthony S. Piro, Sarah Luoma, Arun K. Haldar, Seungmin Hwang, Jennifer Martinez, Meta J. Kuehn, Gregory A. Taylor, Jörn Coers

## Abstract

Inflammation associated with gram-negative bacterial infections is often instigated by the bacterial cell wall component lipopolysaccharide (LPS). LPS-induced inflammation and resulting life-threatening sepsis are mediated by the two distinct LPS receptors TLR4 and caspase-4. Whereas the regulation of TLR4 activation by extracellular and phago-endosomal LPS has been studied in great detail, auxiliary host factors that specifically modulate recognition of cytosolic LPS by caspase-4 are largely unknown. This study identifies dynamin-related membrane remodeling proteins belonging to the family of Immunity related GTPases M clade (IRGM) as negative regulators of caspase-4 activation in macrophages. Phagocytes lacking expression of mouse isoform Irgm2 aberrantly activate caspase-4-dependent inflammatory responses when exposed to extracellular LPS, bacterial outer membrane vesicles or gram-negative bacteria. Consequently, Irgm2-deficient mice display increased susceptibility to caspase-4-mediated septic shock *in vivo.* This *Irgm2* phenotype is partly reversed by the simultaneous genetic deletion of the two additional *Irgm* paralogs *Irgm1* and *Irgm3,* indicating that dysregulated Irgm isoform expression disrupts intracellular LPS processing pathways that limit LPS availability for caspase-4 activation.

## INTRODUCTION

Severe sepsis is an overwhelming microbe-induced inflammatory response resulting in organ dysfunction and constitutes a major public health concern due to high mortality rates and lack of treatment options (Polat, Ugan et al., 2017). A chief sepsis-inducing agent is lipopolysaccharide (LPS, or endotoxin), an abundant building block of the cell wall of all gram-negative bacteria (Munford, 2008, Simpson & Trent, 2019). Extracellular and endocytosed LPS is sensed by the transmembrane receptor toll-like receptor 4 (TLR4), which signals to induce the expression of pro-inflammatory cytokines such as tumor necrosis factor alpha (TNFα) and pro-interleukin-1beta (pro-IL-1β) (Rosadini & Kagan, 2017). Cytosolic LPS on the other hand is sensed by the cysteine protease caspase-4 (originally named caspase-11 in mouse), which upon direct binding to LPS oligomerizes and proteolytically activates the cell death-inducing protein gasdermin D (GSDMD) (Hagar, Powell et al., 2013, Kayagaki, Wong et al., 2013, Liu, Zhang et al., 2016, Shi, Zhao et al., 2015, Shi, Zhao et al., 2014). GSDMD forms pores in the plasma membrane, causing pyroptosis directly as well as indirectly through potassium efflux-mediated activation of the canonical NLRP3 inflammasome (Liu et al., 2016, Ruhl & Broz, 2015, Shi et al., 2015). Activated caspase-1, the defining component of canonical inflammasomes, cleaves numerous substrates including pro-IL-1β to produce mature IL-1β for secretion (Brewer, Brubaker et al., 2019, Broz & Dixit, 2016).

Whereas regulation of TLR4 signaling has been extensively studied, mechanisms that prevent excessive caspase-4-induced inflammation are just beginning to be understood (Rathinam, Zhao et al., 2019). The host-derived oxidized phospholipid oxPAPC as well as stearoyl lysophosphatidylcholine competitively block binding of LPS to caspase-4 (Chu, Indramohan et al., 2018, Li, Zhang et al., 2018). However, oxPAPC was also shown to function as a caspase-4 agonist in dendritic cells and skews LPS-primed macrophages towards proinflammatory metabolism (Di Gioia, Spreafico et al., 2020, Zanoni, Tan et al., 2016, Zanoni, Tan et al., 2017). Additional known checkpoints on caspase-4-triggered inflammation operate downstream from cytosolic LPS sensing and involve inhibition of caspase oligomerization, reduction in membrane lipid peroxidation and membrane repair by the ESCRT III pathway (Choi, Kim et al., 2019, Kang, Zeng et al., 2018, Ruhl, Shkarina et al., 2018).

Similar to the limited number of studies exploring regulatory mechanisms that operate downstream from cytosolic LPS sensing, investigators only recently began to characterize the upstream pathways that control LPS accessibility to the cytosolic compartment (Rathinam et al., 2019). Phagocytosis of bacterial LPS-replete outer membrane vesicles (OMVs) triggers potent activation of caspase-4 (Vanaja, Russo et al., 2016). The mechanism by which LPS derived from endocytosed OMVs escapes into the cytosol to be detected by caspase-4 is not understood but depends on the function of interferon (IFN)-inducible host guanylate binding proteins (Gbps) (Finethy, Luoma et al., 2017). Gbps are additionally required for fast-kinetics caspase-4 activation following gram-negative bacterial infections or cell transfection with purified LPS (Finethy, Jorgensen et al., 2015, Lagrange, Benaoudia et al., 2018, Meunier, Dick et al., 2014, Pilla, Hagar et al., 2014), the latter result implying an additional role for Gbps in caspase-4 activation downstream from LPS release into the host cell cytosol. Uptake of circulating ‘free’ plasma LPS *in vivo* is largely mediated by the hepatocyte-secreted high mobility group box 1 (HMGB1) protein. LPS bound to HMGB1 is endocytosed by macrophages via the receptor for advanced glycation end products (RAGE) and subsequently released into the host cell cytosol through destabilization of LPS-containing endosomes (Deng, Tang et al., 2018). Whether and how the host could negatively regulate any of these different LPS uptake and delivery mechanisms is currently unknown but such regulatory mechanisms would likely require the function of membrane remodeling proteins.

The Immunity related GTPases (IRGs) are dynamin-like proteins that bind to various intracellular membrane compartments, where they ostensibly modulate membrane dynamics and trafficking. IRGs are produced in response to IFNs as well as other immune stimuli including LPS-TLR4 signaling, and mediate several immune programs in response to those agents (Hunn, Feng et al., 2011, Pilla-Moffett, Barber et al., 2016). Variants of the human IRG M clade encoding *IRGM* gene are associated with increased risk for Crohn’s Disease, as first demonstrated by two large genome wide association studies (Parkes, Barrett et al., 2007, Weersma, Stokkers et al., 2009). Subsequent candidate gene association studies expanded the association of human *IRGM* variants to increased susceptibility to mycobacterial infection (Intemann, Thye et al., 2009, King, Lew et al., 2011), and most pertinently, poor outcomes from sepsis (Kimura, Watanabe et al., 2014). While these studies linked the *IRGM* gene to inflammatory diseases, the cellular mechanism by which IRGM suppresses inflammation during infection and the causative genetic alteration in human *IRGM* disease-associated alleles remain poorly defined.

The human genome encodes a single *IRGM* gene, which expresses at least 5 distinct hIRGM splice isoforms. The mouse genome on the other hand encodes 3 paralogous *Irgm* genes, named *Irgm1, Irgm2,* and *Irgm3* (Bekpen, Hunn et al., 2005). The exact functional relationship between hIRGM isoforms and their mouse Irgm orthologs is currently not well defined. However, several recent studies demonstrated that hIRGM and mouse Irgm1 commonly regulate cellular processes that include mitochondrial dynamics (Henry, Schmidt et al., 2014, Liu, Gulati et al., 2013, Schmidt, Fee et al., 2017, Singh, Ornatowski et al., 2010), autophagic flux (Gutierrez, Master et al., 2004, Singh, Davis et al., 2006, Traver, Henry et al., 2011) and NLRP3 inflammasome activation (Mehto, Jena et al., 2019). The parallels between human and mouse IRGMs extend beyond these cell culture phenotypes: similar to hIRGM, Irgm1 attenuates intestinal inflammation (Liu et al., 2013, Rogala, Schoenborn et al., 2018) and promotes host survival during LPS-induced sepsis (Bafica, Feng et al., 2007). Increased mortality among endotoxemic *Irgm1*-/- mice is largely due to hyperresponsive TLR4 signaling in these animals, which leads to augmented TNFα production and inflammation (Bafica et al., 2007, Schmidt et al., 2017). Whereas the anti-inflammatory function of Irgm1 is now firmly established, possible immune-modulatory functions of the two other Irgm paralogs have remained unexplored.

In this study, we systematically assessed the role of individual Irgm proteins in shaping endotoxemic inflammation and discovered a novel function for Irgm2 as an inhibitor of LPS-triggered caspase-4 activation. Accordingly, Irgm2-deficient mice display caspase-4-dependent elevated IL-1β serum levels and more readily succumb to LPS-induced sepsis. Unexpectedly, the simultaneous deletion of all Irgm paralogs partially restores immune homeostasis, thus revealing inter-regulatory relationships among Irgm proteins important for balancing inflammation during gram-negative infections.

## RESULTS

### Irgm1 and Irgm2 but not Irgm3 limit the production of inflammatory cytokines in response to LPS *in vivo*

Mice deficient for *Irgm1* and *Irgm3* were previously reported (Collazo, Yap et al., 2001, Taylor, Collazo et al., 2000). To systematically dissect the physiological function of all three mouse Irgm isoforms, we generated an *Irgm2*-deficient mouse strain. Lack of Irgm2 protein expression was confirmed by Western blotting and immunofluorescence (Appendix Fig S1A-B). To profile the *in vivo* function of individual Irgm isoforms during endotoxemia, we injected wild-type C57BL/6J (WT), *Irgm1-/-, Irgm2-/*-, and *Irgm3-/-* mice with LPS at a concentration of 8 mg/kg bodyweight, collected serum at 4 hours post-injection (hpi) and performed multiplex quantitative serum cytokine measurements. Whereas the LPS-induced cytokine profile of *Irgm3-/-* mice was comparable to WT mice, serum concentrations of several cytokines were elevated in *Irgm1-/-* and *Irgm2-/-* relative to WT (Figs 1A and EV1). Consistent with the previously reported role for Irgm1 as an attenuator of TLR4 signaling (Bafica et al., 2007), we detected significantly elevated TNFα serum concentrations in LPS-treated *Irgm1-/-* mice. Notably, *Irgm2-/-* mice exhibited WT-like TNFα serum levels but increased serum levels for the inflammasome-dependent cytokines IL-1α and IL-1β (Figs 1A and EV1). To validate our multiplex data and to further asses a potential role for Irgm2 in regulating inflammasome-dependent cytokine production *in vivo,* we injected WT, *Irgm1-/-*, *Irgm2-/-* and *Casp1-/-Casp4-/-* mice with LPS and assayed serum IL-18, an additional inflammasome-processed cytokine, as well as IL-1β and TNFα concentrations via ELISA. We detected elevated concentration of IL-1β and IL-18 but not of TNFα in the serum of LPS-injected *Irgm2-/-* mice by ELISA (Fig 1B), confirming our initial observations and thus implicating Irgm2 in the suppression of LPS-induced inflammasome activation *in vivo*.

**Figure 1.**
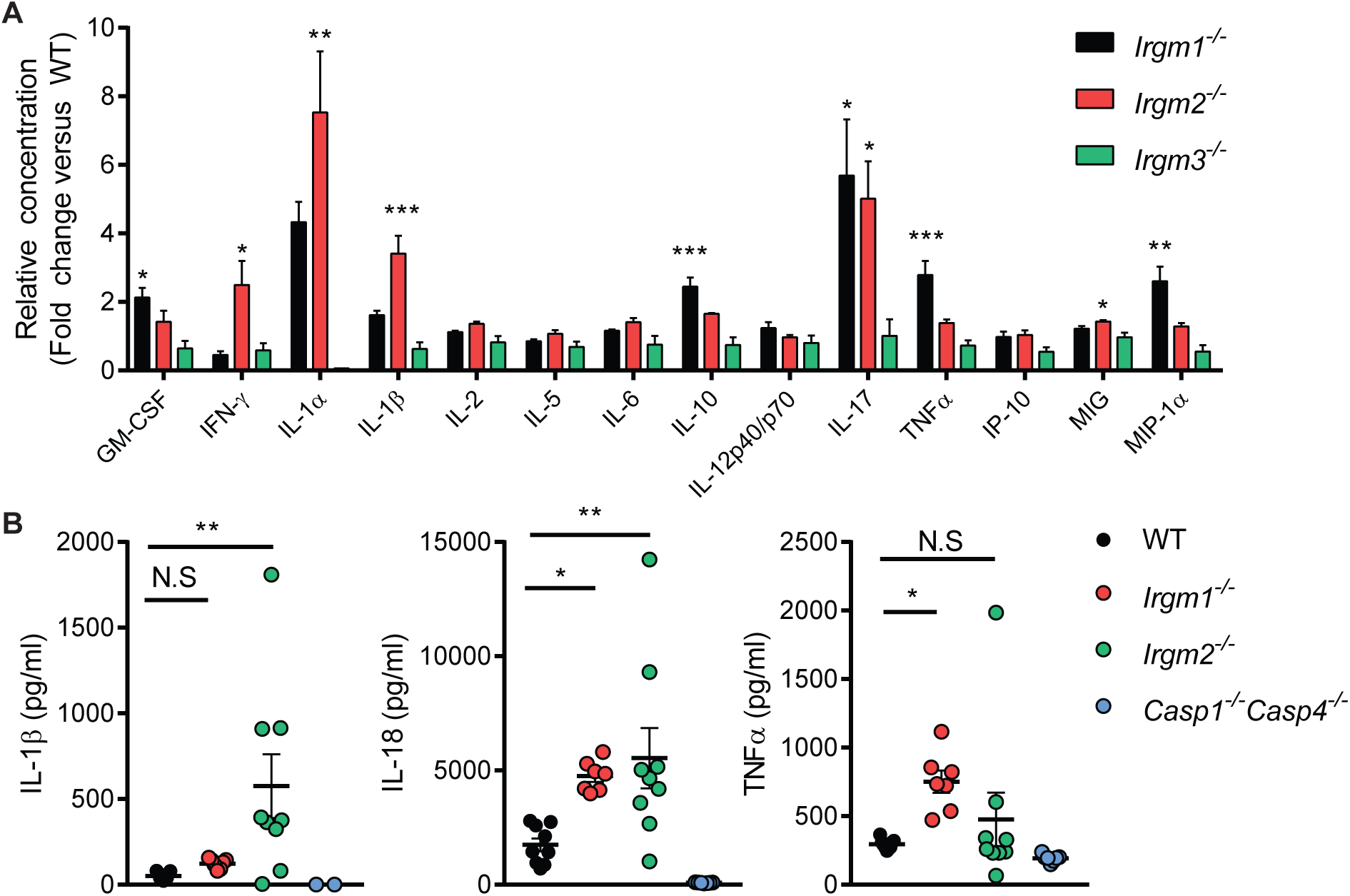
Irgm1 and Irgm2 but not Irgm3 limit the production of inflammatory cytokines in response to LPS *in vivo*. **(A)** WT, *Irgm1*-/-, *Irgm2*-/-*and Irgm3*-/- mice (n = 4 mice/genotype) were injected *i.p.* with LPS (8 mg/kg). Serum was collected 4 hours post injection (hpi) and concentration of various cytokines determined via a preconfigured Luminex multiplex panel. Relative concentration (fold change relative mean of WT) is shown for the indicated cytokines (absolute cytokine concentrations of same experiment are shown in Fig EV1). **(B)** WT (n = 9), *Irgm1*-/- (n = 7), *Irgm2*-/- (n = 9) *and Casp1*-/-*Casp4*-/- (n = 7) mice were injected *i.p.* with LPS (8 mg/kg). Serum was collected 4 hpi and concentration of IL-1β, IL-18, and TNFα was measured via ELISA. Data information: Data shown are means ± SEM. *p < 0.05, **p < 0.01, ***p < for indicated comparisons or comparison between wildtype and indicated genotype by one-way ANOVA with Dunnett’s multiple comparison test. N.S, non significant.

### Irgm2 suppresses inflammasome activation in LPS-treated macrophages

To define the mechanism by which Irgm2 regulates cytokine production, we employed bone marrow-derived macrophages (BMMs) as an established tissue culture model to study LPS-dependent IL-1β secretion. Inflammasome activation in BMMs and consequential secretion of IL-1β is dependent on two signals: the first signal is provided by extracellular damage- or pathogen-associated molecular patterns (DAMPs or PAMPs), such as LPS, that induce the expression of inflammasome components and cytokine proforms; cytoplasmic PAMPs or DAMPs act as a second signal to trigger inflammasome assembly (Broz & Dixit, 2016). To provide the first signal, we primed WT, *Irgm2*-/- and C*asp1-/- Casp4-/-* BMMs with LPS. In all genotypes, LPS priming induced WT levels of pro-IL-1β and TNFα mRNA, and also induced WT-like TNFα secretion, indicating that TLR4 signaling is unchanged in *Irgm2*-/- BMMs (Figs 2A-B). Unexpectedly, 24 hour priming with extracellular LPS at 1 µg/ml was sufficient to induce robust secretion of IL-1β and IL-18 in *Irgm2*-/- but not WT BMMs (Fig. 2B). We considered the hypothesis that *Irgm2*-/- BMMs no longer required a second signal and would therefore secrete IL-1β in response to a broad range of first signals. However, we found that among several tested TLR agonists only LPS was sufficient to trigger IL-1β and IL-18 secretion in *Irgm2*-/- BMMs (Fig 2C), suggesting that extracellular LPS, in the absence of Irgm2 expression, serves as both the first and second signal to activate inflammasomes.

**Figure 2.**
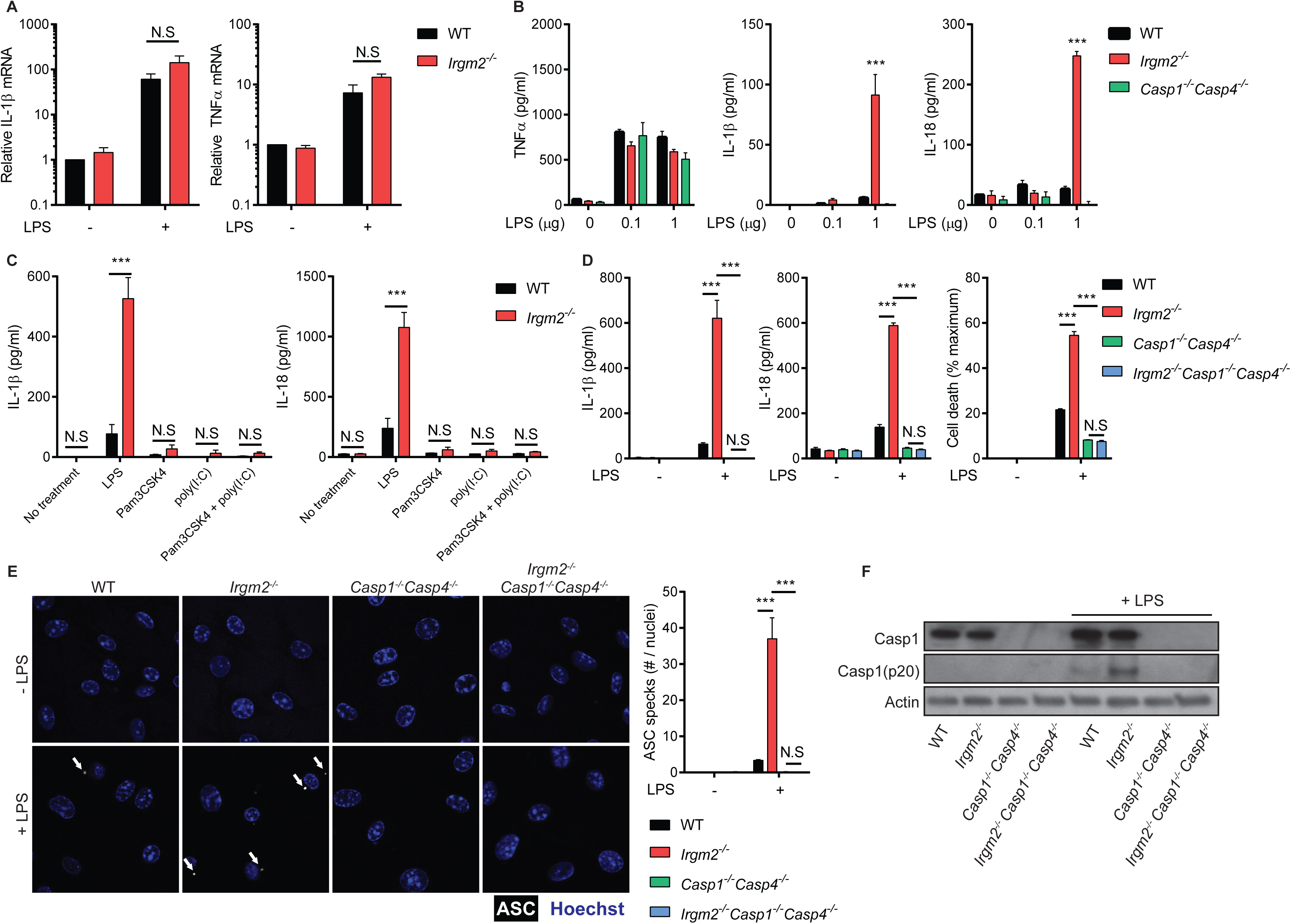
Irgm2 suppresses inflammasome activation in LPS-treated macrophages. **(A)** qPCR measurement of IL-1β, and TNF-α mRNA levels in WT and *Irgm2*-/- BMMs following 8-hour stimulation with LPS (1 µg/ml). **(B)** WT, *Irgm2*-/- and *Casp1*-/-*Casp4*-/- BMMs were treated for 24 hours with LPS at indicated doses and supernatant TNFα, IL-1β, and IL-18 was measured by ELISA. **(C)** IFNγ-primed WT and *Irgm2*-/- BMMs were treated with LPS, Pam3CSK4, poly(I:C) or a combination of Pam3CSK4 and poly(I:C) (1 µg/ml for all treatments) for 24 hours and cell supernatant IL-1β and IL-18 concentrations were assessed by ELISA. **(D)** WT, *Irgm2*-/-, *Casp1*-/-*Casp4*-/- and *Irgm2*-/-*Casp1*-/-*Casp4*-/- BMMs were treated with LPS (1 µg/ml) and IL-1β, IL-18, and LDH release were assessed at 24 hours post treatment (hpt). **(E)** IFNγ-primed WT, *Irgm2*-/-, *Casp1*-/-*Casp4*-/- and *Irgm2*-/-*Casp1*-/-*Casp4*-/- BMMs were treated with LPS (5 µg/ml) for 4 hours and subsequently stained with anti-ASC antibody and Hoechst stain (DNA/nuclei). Representative images are shown with white arrows pointing at ASC specks. Number of ASC specks per nuclei was quantified. **(F)** IFNγ-primed WT, *Irgm2*-/-, *Casp1*-/-*Casp4*-/- and *Irgm2*-/-*Casp1*-/-*Casp4*-/- BMMs were treated with LPS (1 µg/ml) for 24 h and cell lysates and supernatants collected. Protein levels in cell lysates (caspase-1, and actin) and supernatants (caspase-1 p20) were visualized via immunoblotting. Data information: Graphs in (A-E) show means ± SEM from n = 3 independent experiments. ***p < 0.001 for indicated comparisons or comparison between wildtype and indicated genotype by two-way ANOVA with Sidak’s (A, C) or Tukey’s (B, D, E) multiple comparisons test. Images in panel E are representative of one of three independent experiments. Panel F is representative of n = 2 independent experiments. N.S, non significant.

To further test the hypothesis that Irgm2 blocks inflammasome-dependent cellular responses, we monitored pyroptotic cell death in LPS-primed cells. In congruence with our cytokine data, LPS priming resulted in significantly higher rates of cell death in *Irgm2*-/- compared to WT BMMs (Fig 2D). Ectopic expression of Irgm2 in *Irgm2*-/- immortalized BMMs (iBMMs) restored cell viability to WT iBMMs levels (Appendix Fig S2A-B), ruling out the possibility that the death phenotype is due to bystander mutations associated with the *Irgm2* loss-of-function allele. Deletion of the inflammatory caspases caspase-1 and caspase-4 restored cell viability and abolished IL-1β/IL-18 secretion in LPS-primed *Irgm2*-/- BMMs (Fig 2D), indicating that Irgm2 expression suppresses canonical inflammasome activation by extracellularly supplied LPS. To directly visualize canonical inflammasome formation in BMMs, we stained cells for the canonical inflammasome adapter protein Apoptosis-associated Speck-like protein containing a CARD (ASC). During canonical inflammasome assembly ASC polymerizes into a large bioactive protein complex that is visible as a single large speck per individual cell (Broz & Dixit, 2016). Consistent with our predictions, loss of Irgm2 enhanced LPS-induced ASC speck formation (Fig 2E) and increased the amount of autoproteolytically activated caspase-1 p20 (Fig 2F). Collectively, these observations reveal a novel role for Irgm2 in mitigating inflammasome responses.

### Irgm2 suppresses caspase-4 activation in response to LPS and bacterial infections

Because LPS alone is sufficient to trigger inflammasome-mediated cellular responses in *Irgm2*-/- BMMs, we hypothesized that the IFN-inducible cytosolic LPS sensor caspase-4 was activated by extracellularly supplied LPS in cells deficient for Irgm2. In support of this hypothesis, we noticed that IFNγ priming accelerated the kinetics of pyroptotic cell death in *Irgm2*-/- BMMs (Appendix S3A-B). However, to test our hypothesis directly, we monitored inflammasome-dependent responses in BMMs obtained from newly cross-bred *Irgm2-/-Casp4 -/-* mice. Deletion of caspase-4 in *Irgm2*-/- BMMs repressed ASC foci formation, restored cell viability and abrogated LPS-elicited IL-1β/IL-18 secretion (Fig 3A-B). Together, these data show that extracellularly supplied LPS is able to robustly activate the cytosolic LPS sensor caspase-4 when cells lack Irgm2 expression.

**Figure 3.**
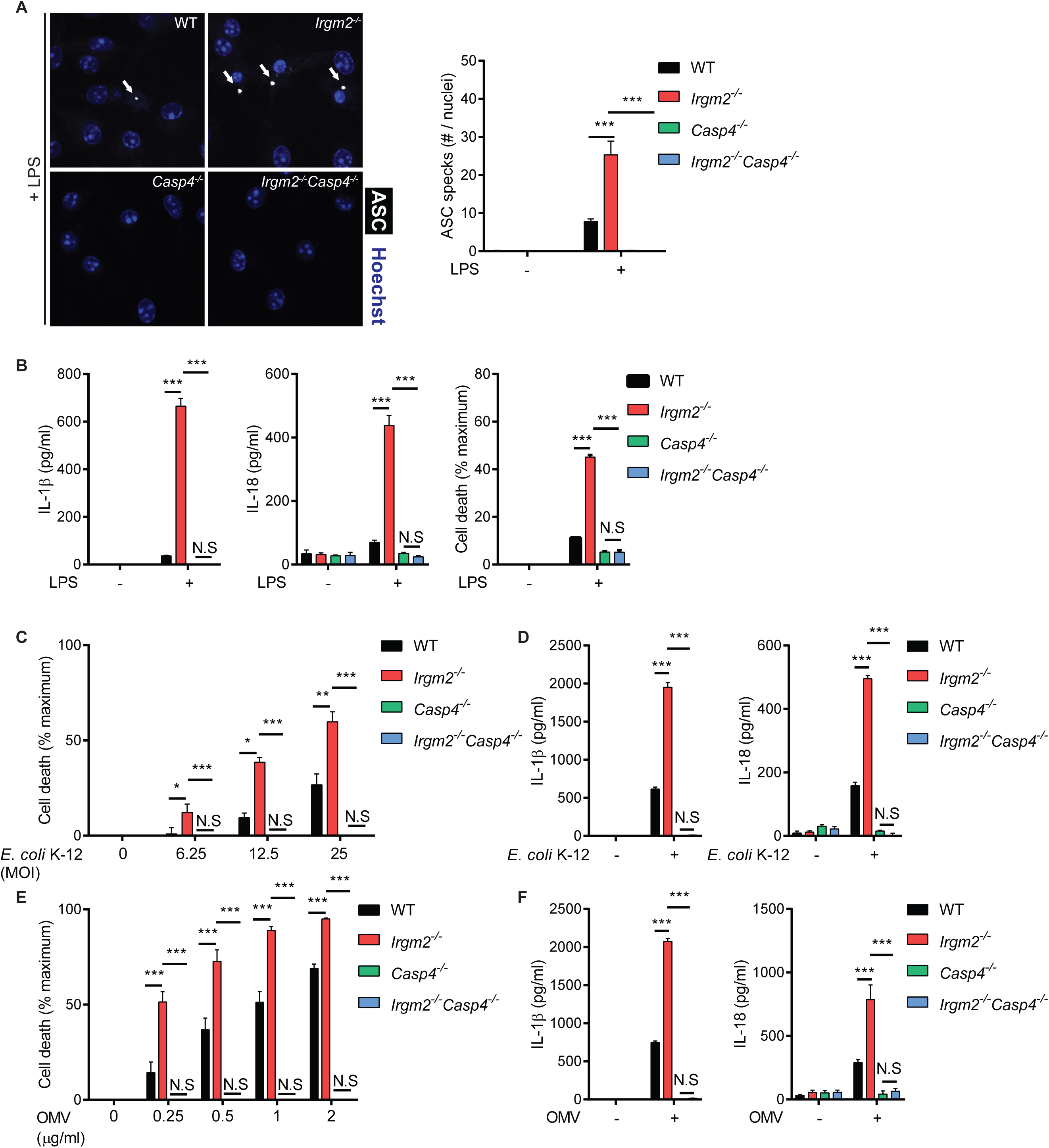
Irgm2 suppresses caspase-4 activation in response to LPS and bacterial infections. **(A)** IFNγ primed WT, *Irgm2*-/-, *Casp4*-/- and *Irgm2*-/- *Casp4*-/- BMMs were treated with LPS (5 µg/ml) for 4 h. Following treatment, cells were stained with anti-ASC antibody and Hoechst (DNA/nuclei). Representative images of ASC specks (white arrows point at specks) are shown and number of ASC specks per nuclei quantified. **(B)** WT, *Irgm2*-/-, *Casp4*-/- and *Irgm2*-/- *Casp4*-/- BMMs were treated with LPS (1 µg/ml). IL-1β, IL-18, and LDH release were assessed at 24 hours-post treatment (hpt). **(C)** WT, *Irgm2*-/-, *Casp4*-/- and *Irgm2*-/- *Casp4*-/- BMMs were infected with *E. coli* K-12 at indicated MOIs and cell viability assessed via CellTiter-Glo. Cell death was calculated as a function of relative viability to uninfected cells. **(D)** WT, *Irgm2*-/-, *Casp4*-/- and *Irgm2*-/- *Casp4*-/- BMMs were infected with *E. coli* K- 12 (MOI 25) and 24 hpi supernatant IL-1β and IL-18 levels were measured by ELISA. **(E)** WT, *Irgm2*-/-, *Casp4*-/- and *Irgm2*-/- *Casp4*-/- BMMs were treated with OMVs at indicated concentrations for 24 h. Cell viability was assessed via CellTiter-Glo and cell death was calculated as a function of relative viability to untreated cells. **(F)** WT, *Irgm2*-/-, *Casp4*-/- and *Irgm2*-/- *Casp4*-/- BMMs were treated with OMVs (1 µg/ml) and 24 hpt cell supernatant IL-1β and IL-18 levels were measured via ELISA. Data information: Shown are means ± SEM from n = 4 (A) or n = 3 (B-F) independent experiments. *p < 0.05, **p < 0.01, ***p < 0.001 for indicated comparisons by two-way ANOVA with Tukey’s multiple comparisons test. Images in panel A are representative of one of four independent experiments. N.S, non significant.

To determine whether Irgm2-mediated regulation of caspase-4 was unique to purified LPS or extended to other caspase-4-activating stimuli, we tested whether Irgm2 could regulate the inflammatory response to bacterial infections. We confirmed our previous observations (Finethy et al., 2017) that nonvirulent *Escherichia coli* K12 induces caspase-4 activation in BMMs in a dose-dependent manner, and found that K12-elcited responses were significantly exacerbated in *Irgm2*-/- BMMs (Fig 3C-D). Similarly, we found that Irgm2 lessened caspase-4 activation in BMMs infected with virulent *E. coli* species, including adherent invasive E. coli (AIEC) and uropathogenic *E. coli* (UPEC) (Fig EV2), demonstrating that Irgm2 attenuates caspase-4 activation in response to both virulent and avirulent *E. coli*.

Irgm proteins regulate the subcellular localization and function of members of a second family of IFN-inducible dynamin-like GTPase, the Gbps (Haldar, Saka et al., 2013, Traver et al., 2011). Gbps colocalize with phagocytosed bacteria expressing virulence-inducing protein secretion systems (Coers, 2017, Feeley, Pilla-Moffett et al., 2017), promote the bacteriolytic destruction of these targeted virulent bacteria, and thereby release PAMPs from lysed bacteria resulting in the activation of cytosolic pattern recognition receptors (Fisch, Bando et al., 2019, Liu, Sarhan et al., 2018, Man, Karki et al., 2015, Meunier, Wallet et al., 2015). Although Gbps fail to colocalize with avirulent *E. coli,* Gbps still promote caspase-4 activation in BMMs exposed to non-pathogenic bacteria through a process that is largely facilitated by the Gbp-mediated recognition and processing of bacterially secreted OMVs (Finethy et al., 2017, Vanaja et al., 2016). We therefore asked whether Irgm2 could regulate the inflammatory response to OMVs. As with our infection experiments, we observed that loss of Irgm2 expression resulted in higher rates of cell death across a range of OMV concentrations (Fig 3E), and similarly amplified OMV-stimulated IL-1β and IL-18 secretion (Fig 3F). These data thus define Irgm2 as an inhibitor of LPS-, OMV- and infection-induced caspase-4 activation.

### Irgm2 regulates caspase-4 activity in a Gbp-independent manner

In light of the functional interactions that exist between Gbp and Irgm proteins (Coers, Brown et al., 2018), we considered the hypothesis that Irgm2 could attenuate inflammasome activation by acting as a Gbp antagonist. We first monitored expression of Gbp2, a protein required for activation of both the noncanonical caspase-4 and the canonical absent in melanoma 2 (AIM2) inflammasomes (Gomes, Cerqueira et al., 2019, Man, Place et al., 2017). Gbp2 as well as caspase-4 were expressed normally in *Irgm2*-/- BMMs (Fig 4A). To functionally test whether Irgm2 modulates Gbp2-dependent activation of AIM2, we infected WT, *Aim2*-/-, *Gbp*chr3-/- (lacking expression of *Gbp1*, *Gbp2*, *Gbp3*, *Gbp5* and *Gbp7*) and *Irgm2*-/- BMMs with the cytosol-invading gram-negative bacterium *Francisella novicida.* We observed that *F. novicida* induced AIM2- mediated cell death in a Gbp-dependent manner (Fig 4B), thus supporting previous findings that Gbp-induced bacteriolysis liberates bacterial DNA for recognition by AIM2 (Man et al., 2015, Meunier et al., 2015). Whereas *F. novicida*-induced pyroptosis was reduced in *Gbp*chr3-/- BMMs, cell death remained unchanged in *Irgm2*-/- BMMs (Fig 4B), demonstrating that the absence of Irgm2 leaves Gbp-dependent AIM2 activation unchanged. Based on a previous report describing Gbp5 as a potential regulator of the NLRP3 inflammasome (Shenoy, Wellington et al., 2012), we also asked whether lack of Irgm2 expression directly impacted NLRP3 activation. We found that treatment of cells with the NLPR3 activator nigericin induced the secretion of equivalent amounts of IL-1β and prompted comparable levels of cell death in WT and *Irgm2*-/- BMMs (Fig EV3). Together, these data rejected a model in which Irgm2 acts as a universal antagonist of Gbps. Because the NLRP3 inflammasome mediates pro-IL-1β processing and IL-1β secretion downstream from LPS-induced caspase-4 activation (Broz & Dixit, 2016), these data further suggested that Irgm2 operates upstream of Gbp/caspase-4- depdendent NLRP3 activation.

**Figure 4.**
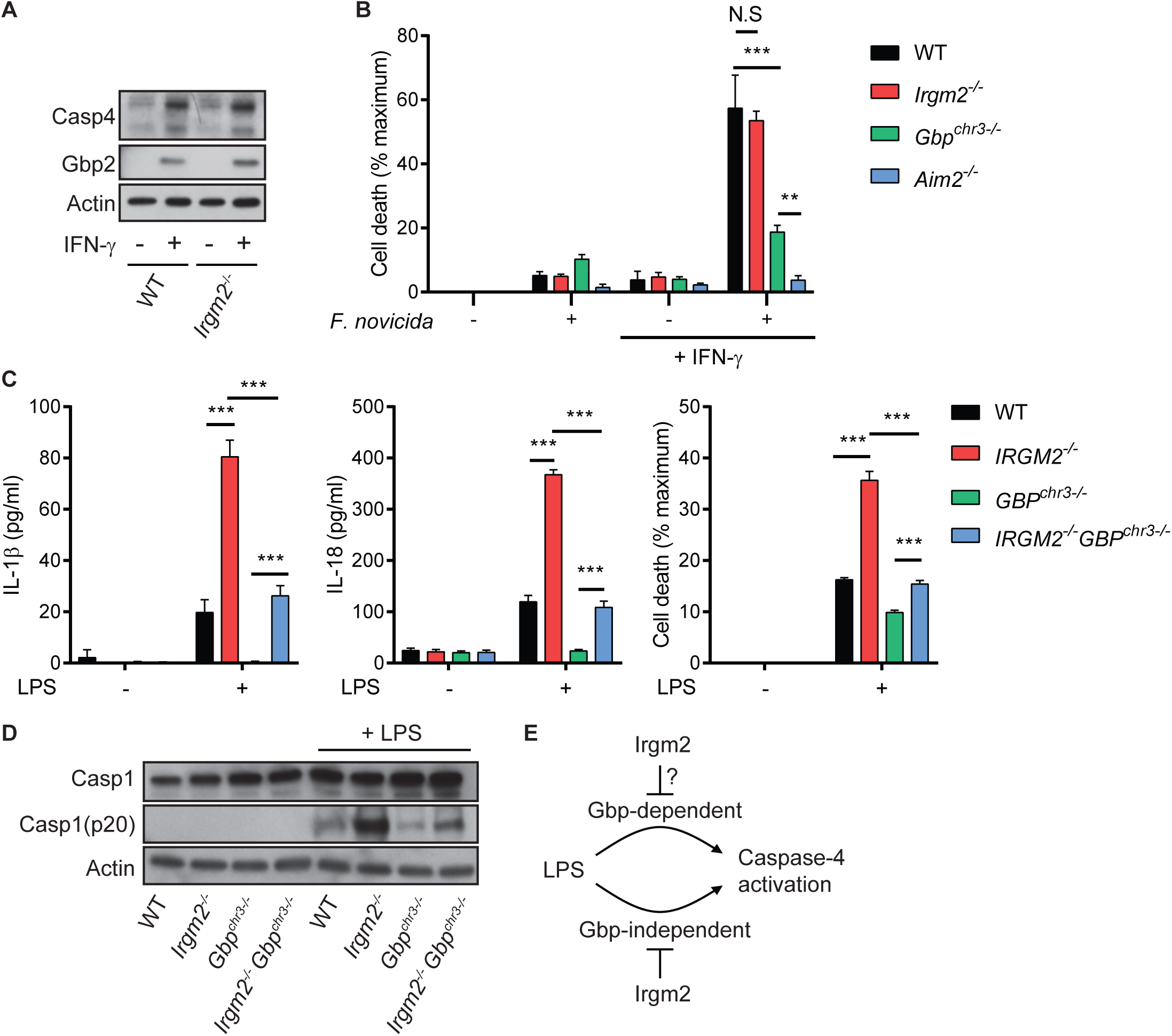
Irgm2 regulates caspase-4 activity in a Gbp-independent manner. **(A)** WT, and *Irgm2*-/- BMMs were stimulated overnight with IFNγ or left untreated and cell lysates were collected. Lysates were assessed for Casp4, Gbp2, and actin protein levels via immunoblotting. **(B)** IFNγ-primed and unprimed WT, *Irgm2*-/-, *Gbpchr3*-/- and *Aim2-/-* BMMs were infected with *Francisella novicida* (MOI 10) and LDH release measured at 4 hpi. **(C)** WT, *Irgm2*-/-, *Gbpchr3*-/- and *Irgm2*-/- *Gbpchr3*-/- BMMs were treated with LPS (1 µg/ml). IL-1β, IL-18, and LDH release were assessed at 24 hpt. **(D)** IFNγ-primed WT, *Irgm2*-/-, *Gbpchr3*-/- and *Irgm2*-/- *Gbpchr3*-/- BMMs were treated with LPS (1 µg/ml) for 24 hours and cell lysates and supernatants collected. Protein levels in cell lysates (Caspase-1, and actin) and supernatants (Caspase-1 p20) were visualized via immunoblotting. **(E)** Model depicting regulation of caspase-4 activation by Irgm2 and Gbps. Data information: Graphs show means ± SEM from n = 3 (B, C) independent experiments. **p < 0.01, ***p < 0.001 for indicated comparisons by two-way ANOVA with Tukey’s multiple comparisons test. (A) and (D) represent one of two independent experiments. N.S, non significant.

We next considered the hypothesis that Irgm2 more narrowly regulates Gbp functions specifically tailored towards the activation of caspase-4. To address this hypothesis, we monitored caspase-4-dependent activation in BMMs taken from newly cross-bred *Irgm2*-/-*Gbp*chr3-/- mice. We found that loss of Irgm2 on a *Gbpchr3-/-* background led to an increase in cell death, enhanced caspase-1 processing and elevated IL-1β/IL- 18 secretion (Fig 4C-D), albeit not to the same level as observed in *Irgm2*-/- BMMs.

While these data do not necessarily allow us to rule out a potential role for Irgm2 in modulating the GBP-dependent caspase-4 pathway, we can conclude that Irgm2 suppresses an additional GBP-independent caspase-4 activation pathway (Fig 4E).

### Irgm2 controls caspase-4 activation upstream from cytosolic LPS sensing

Our results so far proved conclusively that Irgm2 dampens caspase-4 activation in different experimental settings. To determine whether Irgm2 directly regulates cytosolic LPS sensing, we transfected WT, *Irgm2-/-*, *Casp4-/-* and *Irgm2-/-Casp4-/-* BMMs with LPS and measured pyroptotic death. As expected, caspase-4-deficient BMMs were resistant to LPS-mediated cell death, whereas WT BMMs succumbed to cytosolically delivered LPS (Fig EV4A). Remarkably, we found that cell death was equivalent between LPS-transfected WT and *Irgm2-/-* BMMs (Fig EV 4A). To further substantiate these findings, we deposited LPS into the host cell cytosol via co-delivery with the vacuole-disrupting gram-positive bacterium *Listeria monocytogenes.* Using this alternative approach, we again found that WT and *Irgm2-/-* BMMs were equally susceptible to LPS-triggered pyroptotic cell death across a 1-log range of LPS concentrations (Fig EV4B). These data showed that cytosolic delivery of LPS circumnavigates the inhibitory function Irgm2 exerts on caspase-4.

Because Irgm2 blocks caspase-4 activation in response to extracellular but not cytosolic LPS, Irgm2 must prevent extracellular LPS from becoming accessible for cytosolic recognition by caspase-4. However, deletion of an important mediator of LPS uptake, the glycosylphosphatidylinositol (GPI)-anchored LPS-binding protein CD14, did not reduce cell death or IL-18 secretion in LPS-treated *Irgm2-/-* BMMs (Fig EV4C-D).

These data thus suggest that a CD14-independent pathway delivers LPS for caspase-4 sensing in *Irgm2-/-* BMMs.

We next asked whether Irgm2 could exert its anti-inflammatory effect simply through the inhibition of LPS ingestion. We therefore monitored the uptake of fluorescently labeled LPS in both unprimed and IFNγ-primed *Casp4-/-* and *Irgm2-/-Casp4-/-* BMMs, choosing caspase-4-deficient cells to avoid compounding effects resulting from pyroptotic cell death. We found that lack of Irgm2 expression failed to increase LPS ingestion (Fig EV4E-F). These data thus suggested that Irgm2 does not control LPS uptake but rather downstream processing of ingested LPS.

### Autophagy related proteins control caspase-4 activation upstream from cytosolic LPS sensing

The autophagy machinery and specifically autophagy-related protein 8 (Atg8) family member Gamma-aminobutyric acid receptor-associated protein (Gabarap) were previously reported to limit NLRP3 activation, and it was shown that autophagy-deficient compared to WT macrophages secrete elevated levels of IL-1β in response to LPS treatment (Harris, Hartman et al., 2011, Saitoh, Fujita et al., 2008, Zhang, Xu et al., 2013). In agreement with these previous studies we found that LPS-treated autophagy-deficient *Atg5-/-* and *Atg14-/-* BMMs secreted elevated amounts of IL-1β and IL-18, formed ASC specks more frequently and underwent cell death more readily when compared to WT BMMs, thus resembling the response we observed in *Irgm2-/-* BMMs (Fig 5A-B). To address whether direct inhibition of the NLRP3 pathways or enhanced LPS accessibility were primarily responsible for the LPS hypersusceptibility phenotype of *Atg5-/-* and *Atg14-/-* BMMs, we transfected these cells with LPS and monitored cell death. We found that LPS transfection induced comparable levels of cell death in *Atg5-/-, Atg14-/-, Irgm2-/-* and WT BMMs (Fig 5C), suggesting that Atg5, Atg14 and Irgm2 are likewise involved in blocking LPS accessibility to the cytosolic sensor caspase-4 rather than the direct regulation of caspase-4 function itself.

**Figure 5.**
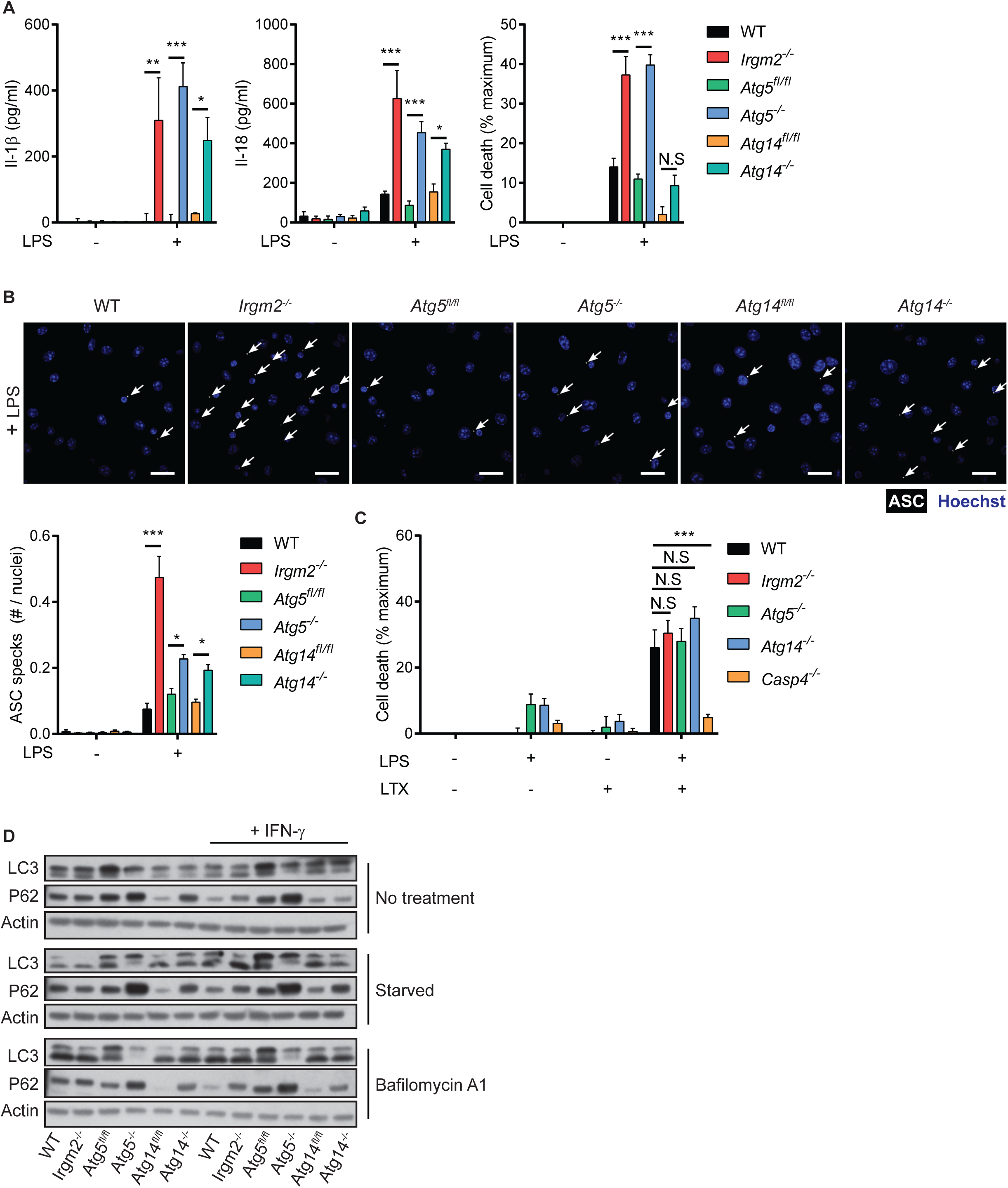
Autophagy related proteins control caspase-4 activation upstream from cytosolic LPS sensing. **(A)** WT, *Irgm2-/-*, *Atg5fl/fl*, LysMCre-*Atg5*f/f (*Atg5*-/-), *Atg14fl/fl* and LysMCre-*Atg14*f/f (*Atg14*-/-) BMMs were treated with LPS (1 µg/ml) and IL-1β, IL-18, and LDH release were assessed at 24 hpt. (n = 4 independent experiments for IL-1β, IL-18 and n = 7 independent experiments for LDH release) **(B)** IFNγ-primed of the indicated genotypes were treated with LPS (5 µg/ml) for 4 hours and subsequently stained with anti-ASC antibody and Hoechst stain (DNA/nuclei). Representative images are shown with white arrows pointing at ASC specks. Number of ASC specks per nuclei was quantified. (n = 3 independent experiments, >200 nuclei counted for each condition/replicate) **(C)** IFNγ-primed WT BMMs of the indicated genotypes were transfected with LPS using lipofectamine LTX and LDH release measured 2 hpt. (n = 5 independent experiments) **(D)** IFNγ-primed WT BMMs of the indicated genotypes were stimulated overnight with IFNγ or left untreated. Cells were then starved for 2 h in HBSS or were treated with Bafilomycin A1 (100 nM) and cell lysates were collected. Lysates were assessed for LC3, p62, and actin protein levels via immunoblotting. Image is representative of n = 3 independent experiments. Data information: Graphs show means ± SEM. *p < 0.05, **p < 0.01, ***p < 0.001 for indicated comparisons by two-way ANOVA with Tukey’s multiple comparisons test. N.S, non significant.

Because *Atg5-/-, Atg14-/-* and *Irgm2-/-* BMMs display comparable phenotypes, we speculated that all of these host factors operate in the same pathway to attenuate caspase-4 responses. In support of this model previous studies had demonstrated that human IRGM as well as mouse Irgm1 modulate autophagosome formation and autophagic flux, at least in part through direct interactions with members of the Atg8 family such as LC3 (Kumar, Jain et al., 2018, Singh et al., 2006). We therefore asked whether the observed LPS sensitization of *Irgm2-/-* BMMs could be the result of an autophagy defect in these cells. To test this hypothesis, we monitored LC3 lipidation and autophagic flux-dependent degradation of the ubiquitin-binding adapter protein p62 in WT, *Irgm2-, Atg5-* and *Atg14*-deficient BMMs. LC3 lipidation and p62 degradation were comparable in WT and *Irgm2-/-* BMMs, essentially excluding a prominent role for Irgm2 in modulating canonical autophagy (Fig 5D), and instead suggesting a possible role for Irgm2 in a non-canonical autophagy-related pathway. This model is supported by findings reported in the accompanying manuscript by Meunier and colleagues, who show that Irgm2 interacts with the Atg8 protein Gabarapl2 to attenuate caspase-4 activation.

### Irgm2 protects against caspase-4-driven septic shock during endotoxemia

Because cytosolic sensing of LPS by caspase-4 promotes septic shock in a mouse model (Hagar et al., 2013, Kayagaki et al., 2013), we hypothesized that loss of Irgm2 would increase susceptibility to an LPS challenge *in vivo*. To test our hypothesis, WT, *Irgm2-/-*, *Casp4-/-* and *Irgm2-/-Casp4-/-* mice were intraperitoneally injected with 8 mg/kg bodyweight LPS and monitored for 48 hours. As predicted, *Irgm2-/-* mice displayed decreased survival rates relative to WT mice and loss of caspase-4 restored survival in Irgm2-deficient and WT animals alike (Fig 6A). Similarly, deletion of caspase-4 completely reduced IL-1β and IL-18 serum levels in LPS-challenged *Irgm2-/-* mice to those detected in *Casp4-/-* mice (Fig. 6B). These findings demonstrate that Irgm2 blocks caspase-4-depenent inflammatory responses initiated by circulating serum LPS.

**Figure 6.**
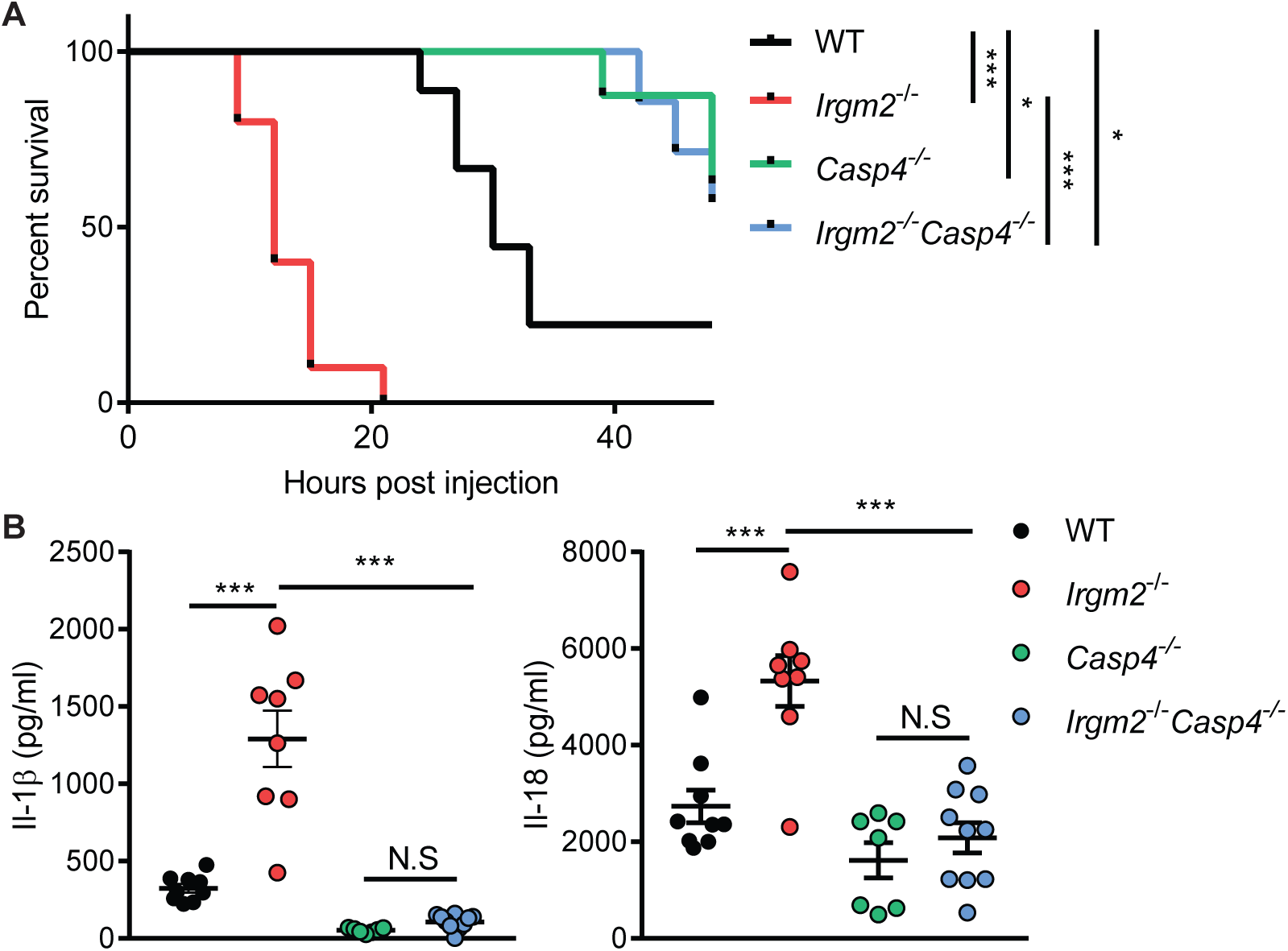
Irgm2 protects against caspase-4-driven septic shock during endotoxemia. **(A)** WT (n = 9), *Irgm2*-/- (n = 10), *Casp4*-/- (n = 7) and *Irgm2*-/-*Casp4*-/- (n = 8) mice were injected *i.p.* with LPS (8 mg/kg). Morbidity and mortality were observed for 48 h at 3 hour intervals. **(B)** WT (n = 9), *Irgm2*-/- (n = 8), *Casp4*-/- (n = 7) and *Irgm2*-/-*Casp4*-/- (n = 10) mice were injected *i.p.* with LPS (8 mg/kg). Serum was collected 4 hpi and concentration of IL-1β, and IL-18 was measured by ELISA. Data shown are from 2 pooled experiments. Data information: Graphs (B) show means ± SEM. *p < 0.05, ***p < 0.001 for indicated comparisons by log-rank Mantel-Cox test (A) or one-way ANOVA with Tukey’s multiple comparisons test (B). N.S, non significant.

### Immune homeostasis is partially restored in mice deficient for all Irgm paralogs

We previously provided evidence in favor of a model in which distinct Irgm isoforms have antagonistic relationships with each other. For example, we showed that T cell homeostasis disrupted in *Irgm1-/-* animals was restored through the simultaneous deletion of Irgm3, a second Irgm family member (Coers, Gondek et al., 2011, Feng, Zheng et al., 2008, Henry, Daniell et al., 2009). To determine whether the disturbance of these inter-regulatory relationships among Irgm protein could contribute to the elevated inflammatory state of LPS-treated *Irgm2-/-* cells and mice, we generated a novel mouse strain deficient for the expression of all 3 Irgm isoforms (Fig EV5A), henceforth referred to as the pan*Irgm-/-* strain. We found that lack of Irgm1 and Irgm3 expression in pan*Irgm-/-* BMMs substantially moderated the inflammatory response of Irgm2-deficient macrophages exposed to extracellular LPS or *E. coli*: specifically, we observed that pan*Irgm-/-* BMMs secreted significantly less IL-1β and were more resistant to pyroptotic cell death than *Irgm2-/-* BMMs (Fig 7A-B). Nonetheless, pan*Irgm-/-* BMMs remained more responsive to LPS and *E. coli* than WT BMMs across different experimental conditions (Fig 7A-B), indicating that the Irgm protein network as a whole exerts an anti-inflammatory effect. In agreement with our cell culture observations, we found that deletion of Irgm1 and Irgm3 partially reversed the caspase-4-driven hyperinflammatory response of Irgm2-deficient animals, as evident by the intermediate IL-1β serum levels of endotoxemic pan*Irgm-/-* relative to WT and *Irgm2-/-* mice (Fig 7C). TNFα serum levels were comparable across the different genotypes tested (Fig 7C), indicating that deletion of all Irgm isoforms not only partially normalized the sensitized caspase-4 pathway of *Irgm2-/-* mice but also tempered the excessive TLR4 signaling pathway of *Irgm1-/-* mice (Bafica et al., 2007, Schmidt et al., 2017). Lastly, we found that the deletion of all *Irgm* genes increased survival of endotoxemic mice to WT or near WT levels (Figs 7D and EV5B), supporting a model, according to which dysregulated Irgm isoform expression rather than merely Irgm2 loss-of-function drives the pathological inflammation in endotoxemic *Irgm2-/-* mice.

**Figure 7.**
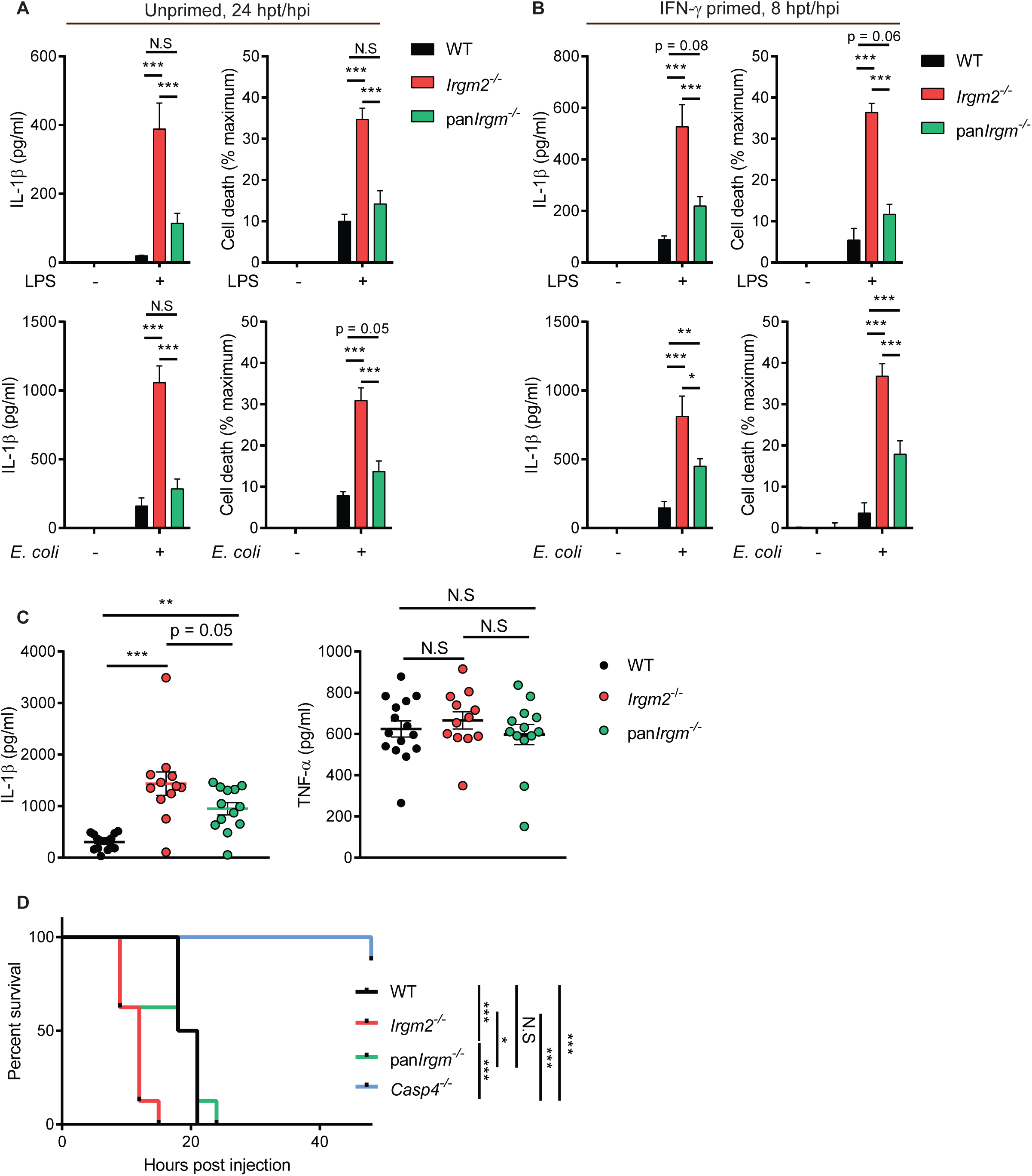
Immune homeostasis is partially restored in mice deficient for all Irgm paralogs. **(A)** WT, *Irgm2*-/-, and pan*Irgm*-/- BMMs were treated with LPS (1 µg/ml) or infected with *E. coli* K-12 (MOI 25) and IL-1β, and LDH release were assessed at 24 hpt. (n = 3 independent experiments for LPS treatments and n = 6 independent experiments for *E. coli* infections) **(B)** IFNγ-primed WT, *Irgm2*-/-, and pan*Irgm*-/- BMMs were treated with LPS (1 µg/ml) or infected with *E. coli* K-12 (MOI 25) and IL-1β, and LDH release were assessed at 8 hpt/ hpi. (n = 3 independent experiments for IL-1β release, n = 5 independent experiments for LDH) **(C)** WT (n = 15), *Irgm2*-/- (n = 12), and pan*Irgm*-/- (n = 13) mice were injected *i.p.* with LPS (8 mg/kg). Serum was collected 4 hpi and concentration of IL-1β, and IL-18 was measured via ELISA. Data shown are from 3 pooled experiments. **(D)** WT, *Irgm2*-/-, pan*Irgm*-/- and *Casp4*-/- mice (n = 8 mice/genotype) were injected *i.p.* with LPS (8 mg/kg body weight). Morbidity and mortality were observed for 48 h at 3-h intervals. Data information: Graphs show means ± SEM. *p < 0.05, **p < 0.01 ***p < 0.001 for indicated comparisons by two-way ANOVA (A, B) or one-way ANOVA (C) with Tukey’s multiple comparison test, or log-rank Mantel-Cox test (D). N.S, non significant.

## DISCUSSION

Infections elicit robust inflammatory responses, which benefit the host by boosting protective antimicrobial immunity. However, dysregulated or overly robust inflammation can cause substantial host morbidity and mortality. Host organisms including humans thus evolved complex regulatory mechanisms to calibrate inflammation so that infections can be contained while collateral damage to infected tissue is minimized (Casadevall & Pirofski, 1999). Here we define a novel pathway responsible for regulating and minimizing inflammation elicited by the cytosolic LPS sensor caspase-4. Our studies characterize mouse Irgm proteins as novel rheostats calibrating the sensitivity of cells and whole animals to LPS. We demonstrate that deletion of Irgm2 leads to excessive activation of the noncanonical caspase-4 inflammasome pathway. We further show that simultaneous deletion of all Irgm paralogs largely restores immune homeostasis in Irgm2-deficient animals, thus indicating that the disturbance of the Irgm expression network is largely responsible for LPS hyperresponsiveness and inflammation in these animals.

While caspase-4 is known to sense LPS in the cytosol, how LPS becomes accessible to its cytosolic sensor is not well understood. The majority of ‘free’ LPS is internalized via receptor-mediated endocytosis or macropinocytosis and then trafficked to the lysosomal compartment for detoxification (Deng et al., 2018, Kitchens, Wang et al., 1998, Poussin, Foti et al., 1998). How LPS exits membrane-bound compartments has remained mostly unstudied owing in part to the observation that when WT macrophages are exposed to extracellular LPS in cell culture, only minimal caspase-4 activation is detectable (Hagar et al., 2013, Kayagaki et al., 2013). Here, we demonstrate that extracellular LPS can potently activate the inflammasome in murine macrophages deficient for Irgm2.

There are a variety of nodes in the LPS uptake and processing pathway, which could potentially be modulated by Irgm2 including i) lysosomal LPS detoxification, ii) LPS internalization, and iii) LPS accessibility. The lysosomal enzyme acyloxyacyl hydrolase (AOAH) deacylates LPS and thereby inactivates LPS as a caspase-4 and TLR4 agonist. Consequently, deletion of AOAH results in prolonged reactions to LPS *in vivo* (Lu, Varley et al., 2013, Lu, Varley et al., 2008). However, AOAH operates with slow kinetics and inactivates only 25% of LPS over a twelve-hour period (Ojogun, Kuang et al., 2009). We therefore predict that even complete loss of AOAH could only have a minor effect on caspase-4 activation in the shorter time courses used in this study. A more likely model would predict a role for Irgm2 in LPS internalization. While Irgm2 has never been associated with internalization pathways, a previous study linked Irgm1 expression to internalization of oxidized low density lipoprotein (Xia, Li et al., 2013). This study reported that Irgm1 promotes CD36-mediated lipoprotein uptake by regulating F-actin polymerization. Irgm2 could play an opposing role and minimize lipid internalization or if, like Irgm1, Irgm2 regulates actin polymerization, loss of Irgm2 could lead to increased macropinocytosis promoting bulk nonspecific uptake of extracellular LPS. Arguing against this model, we found that bulk LPS uptake by macrophages was not noticeably changed in Irgm2-deficient macrophages. As an alternative model, we propose that Irgm2 controls cytosolic accessibility of LPS. As far as we know, no mammalian LPS transmembrane transporter exists. However, a recent report demonstrated that hepatocyte-secreted HMGB1 binds to circulating serum LPS, promotes LPS uptake into lysosomes and facilitates LPS delivery from lysosomes into the host cell cytosol (Deng et al., 2018). Based in part on the observation that amphipathic HMGB1 can destabilize liposomes at an acidic pH *in vitro,* it was proposed that HMGB1 disrupts lysosomal membranes and thereby promotes cytosolic leakage of LPS (Deng et al., 2018). While our cell culture studies were conducted in the absence of HMGB1, it is conceivable that amphipathic LPS itself or other host-derived molecules could induce moderate vacuolar destabilization of LPS-containing compartments, especially in IFN-stimulated or LPS-primed macrophages. For example, radical oxygen production and lipid peroxidation were shown to promote antigen release from endosomes via membrane destabilization (Dingjan, Verboogen et al., 2016, Kotsias, Hoffmann et al., 2013). We predict that Irgm2 minimizes membrane damage of LPS-containing vacuoles and thereby blocks LPS accessibility for caspase-4. The expression of Irgm2 could promote vacuolar stability through a number of potential mechanisms such as blocking reactive oxygen species production, repairing damaged endosomes or sorting LPS-containing endosomes away from rupture-prone lysosomes. Additional studies are necessary to define if and how Irgm2 regulates each of these processes.

While our results reveal a novel function for mouse Irgm2 in controlling caspase-4 activation, it is unclear whether Irgm2 shares this function with human IRGM. Prior studies of mouse Irgm proteins almost exclusively focused on Irgm1 and revealed shared functions of mouse Irgm1 and human IRGM, especially regarding the regulation of mitochondrial dynamics (Coers et al., 2018). Mitochondrial dysfunction and the associated release of oxidized mitochondrial DNA provide compelling molecular models to account for the increased inflammation observed in Irgm1-deficient cells and animals, and may also underlie some of the autoinflammation associated with human *IRGM* disease alleles (Coers et al., 2018, Holley & Schroder, 2020). In addition to controlling mitochondrial remodeling and autophagy, Golgi-resident human IRGM also regulates Golgi fragmentation in response to viral infections (Hansen, Johnsen et al., 2017). Whether mouse Irgm proteins execute similar functions has not been explored, but the localization of both Irgm1 and Irgm2 to the Golgi Apparatus ((Henry et al., 2014, Martens & Howard, 2006) and Fig S1) indicates that murine IRGM homologs may play similar roles in shaping Golgi membrane remodeling events. Accompanying studies by Meunier and colleagues show that the Irgm2-interaction partner Gabarapl2, another Golgi-resident protein (Sagiv, Legesse-Miller et al., 2000), cooperates with Irgm2 in mouse macrophages and blocks caspase-4 activation in both mouse and human macrophages. Thus, the same anti-inflammatory pathway disrupted in *Irgm2-/-* BMMs appears to also operate in human cells. Future studies will be aimed at defining this pathway in detail through the systematic discovery and characterization of the molecular players and their activities, many of which are likely shared between mouse and human macrophages.

Too much or too little inflammation is host detrimental. The mammalian host thus evolved complex regulatory networks, which fine-tune the inflammatory response during an infection. The identity of many of these regulatory circuits and how they function remain to be determined. Here we report the discovery of a novel regulatory pathway, which attenuates cytosolic LPS sensing and moderates LPS-induced inflammation. Future studies will expand our mechanistic understanding of this regulatory pathway and may open up avenues for therapeutic interventions to alleviate disease during gram-negative bacterial infections.

## MATERIALS AND METHODS

### Production of the *Irgm2-/-*mouse line

To generate *Irgm2-/-* mice, we inserted a *loxP* site together with a *neo* marker (*floxneo*) flanked by two flippase recognition target sites (*frt*) before the ATG of *Irgm2* and a *loxP* site after the stop codon of *Irgm2* using backbone vector pEZ-Frt-lox-DT (Addgene plasmid #11736). We placed this cassette between 5’ and 3’ *Irgm2* homology arms. The *Irgm2* targeting construct was transfected into C57BL/6-derived PRX-B6N ES cells (Primogenix). Properly targeted ES cell clones were identified by PCR screening, confirmed by Southern blotting and used for blastocyst injection. The resulting chimeras were crossed with C57BL/6J mice and germline transmission of targeted ES clones was obtained. We performed Southern blot analysis to confirm the proper targeting in F2 mice. These mice were crossed with a germline flippase-expressing strain B6(C3)-Tg(Pgk1-FLPo)10Sykr/J (Wu, Wang et al., 2009) to remove the neomycin cassette. Offspring from this cross harbored a floxed *Irgm2* allele (*Irgm2f*lox). *Irgm2f*lox mice were crossed with the germline-expressing Cre strain B6.FVB-Tg(EIIa-cre)C5379Lmgd/J (Lakso, Pichel et al., 1996) to generate *Irgm2*-/- mice. Mice were genotyped by PCR using primers Irgm2_543 5’- GATCTGAAAGCCTGGACAA-3’ and Irgm2_542 5’- GGGTACACATGGGGTACAGG-3’ which yield a 1.4 kb (WT) and 0.2 kb (*Irgm2*-/-) DNA product.

### Production of the pan*Irgm-/-* mouse line

To generate pan*Irgm-/-* mice, CRISPR sgRNA-Cas9 protein ribonucleoprotein complexes were injected into pronuclear-stage mouse embryos derived from previously reported double-knockout (*Irgm1*-/-*Irgm3*-/-) mice on a C57BL/6J genetic background (Coers et al., 2011, Henry et al., 2009). The following pair of sgRNA were used for the injection: sgRNApan1 5’-GACTGCCGTGAAAGAAGGGG-3’ and sgRNApan4 5’-GGTTGGCTCTATAGGTTTTG-3’. The genomic *Irgm2* locus of pan*Irgm-/-* founder mice was sequenced and a founder line with 925 bp deletion and nonsense mutation was backcrossed to C57BL/6J and then intercrossed to obtained animals homozygous for the deletion alleles of all 3 *Irgm* paralogs, as confirmed by PCR genotyping and Western blotting.

### Additional mouse strains used

WT (C57BL/6J) mice were originally purchased from Jackson Laboratories. *Irgm1-/-*, *Irgm3-/-*, *Atg5*f/f, LysMCre-*Atg5*f/f, *Atg14*f/f, LysMCre-*Atg14*f/f, *Casp1-/-Casp4-/-*, *Casp4-/-, CD14-/-, Aim2-/-, Nlrp3-/-,* and *Gbpchr3-/-* mice were previously described (Choi, Park et al., 2014, Collazo et al., 2001, Inoue, Williams et al., 2012, Kayagaki, Warming et al., 2011, Kuida, Lippke et al., 1995, Moore, Andersson et al., 2000, Rathinam, Jiang et al., 2010, Taylor et al., 2000, Yamamoto, Okuyama et al., 2012, Zhao, Fux et al., 2008). The mouse strains *Irgm2-/-Casp1-/-Casp4-/-*, *Irgm2-/-Casp4-/-*, *Irgm2-/-GBPchr3-/-* and *Irgm2-/- CD14-/-* mice were generated through intercrossing of aforementioned mouse lines.

### Mouse husbandry and *in vivo* experiments

All mouse lines were maintained at animal facilities at Duke University Medical Center. Animal protocols were approved by the Institutional Animal Care and Use Committees at Duke University. Mice were injected intraperitoneally with 2 or 8 mg *E. coli* LPS (O111:B4 LPS; L3024; Sigma)/ kg bodyweight or with PBS control, as indicated. For cytokine monitoring, serum was collected 4 hours after LPS injection. Cytokine concentrations were measured by ELISA or Luminex Multiplex Assay (Invitrogen Mouse 20-plex). To determine host susceptibility to endotoxemia, mice were monitored every 3 h for up to 48 hours following LPS injection. Mice were considered moribund and euthanized when their body weight dropped below 80% of starting weight, or the mice exhibited convulsions, or uncontrollable shivering, or severe ataxia as indicated by lack of righting response. We used approximately equal numbers of male and female mice for experiments.

### Production and culturing of bone marrow derived macrophages (BMMs)

Bone marrow-derived macrophages (BMMs) were generated essentially as previously described (Finethy et al., 2017, Pilla et al., 2014). Briefly, bone marrow was flushed out from mouse femurs and tibia using a 23-gauge needle and then cultured in RPMI 1640, 20% FBS, 2-mercaptoethanol and 14% conditioned media containing macrophage colony-stimulating factor at 37 °C in 5% (vol/vol) CO_2_ in nontissue culture-treated dishes for 5 – 7 days. Differentiated BMMS were harvested, frozen and then thawed again for use in experiments. Immortalized BMMs (iBMMs) were generated through infection with J2 virus as previously described (Pilla et al., 2014).

### Ectopic expression of Irgm2

Irgm2 was inserted into the tetracycline-inducible vector pInducer20 via Gateway recombination (Meerbrey, Hu et al., 2011). To this end, the Irgm2 cDNA was amplified from RNA isloated from IFNγ-stimulated WT BMMs using the primer pair attB1-IRGM2-Forward/attB2-IRGM-Reverse, which add both flanking attB-recombination sites and a 5’-Kozak consensus sequence. The resulting PCR product was inserted into pDONR221 via Gateway BP recombination, followed by Gateway LR recombination into pInducer20. Viral particles were prepared in HEK-293T cells and transduced into iBMMs as previously described (Feeley et al., 2017). Expression of Irgm2 was induced by overnight incubation with cell culture medium containing anhydrotetracycline (aTc, 2 µg/ml). Primers: attB1-Irgm2-Forward, 5’ GGGGACAAGTTTGTACAAAAAAGCAGGCTGCCACCATGGAAGAGGCAGTTGAGTC ACC-3’; attB2-Irgm2-Reverse, 5’-GGGGACCACTTTGTACAAGAAAGCTGGGTCCTAAGGATGAGGAATGGAGAGTCTC AGG-3’

### Bacterial culture and OMV production

*E. coli* K-12 BW5113 (K12), uropathogenic *E. coli* CFT073 (UPEC), and *E. coli* O18:K1:H7 strain C5 (AIEC) were cultured overnight in Luria-Bertani broth (LB) at 37°C with agitation (Baba, Ara et al., 2006, Johnson, Delavari et al., 2001, Welch, Burland et al., 2002). *Francisella novicida* strain U112 (ATTC catalog No. NR-13) was grown overnight in Brain Heart Infusion (BHI) broth shaking at 37°C. For LPS delivery experiments, *Listeria monocytogenes* strain DH-L1039 was grown overnight in BHI at 30°C, as previously reported (Pilla et al., 2014). OMVs were isolated from culture supernatants via ultracentrifugation and ultrafiltration as previously described (Finethy et al., 2017).

### Tissue culture infection, extracellular LPS and OMV treatment protocols

For all infections, bacteria cultures were diluted in cell culture medium to achieve the desired multiplicity of infection (MOI) and then added directly to cells. Cells were then centrifuged at 700 x g for 10 min. Following 1 hour incubation at 37°C, gentamicin was added to wells at a final concentration of 100 µg/ml and cells were assayed at indicated time points. Where indicated, cells were primed with 100 U/ml of mouse IFNγ (EMD Millipore) overnight. Ultrapure *S. minnesota* R595 LPS (List Biologicals) at a concentration of 1 µg/ml was used for all cell culture LPS treatments unless otherwise noted. Where described, cells were treated with 1 µg/ml poly(I:C) (Sigma-Aldrich), 1 µg/ml Pam3CSK4 (Invivogen), or both. *E. coli* K-12 BW25113. To assess cytokine production or cell death, agonists were diluted in complete cell culture medium and added directly to cells for designated times. For qPCR experiments, cells were treated with LPS or left untreated for 8 hours prior to harvesting of RNA. For immunoblot experiments, cells were washed 3 times with PBS before addition of LPS diluted in Opti-MEM (Gibco). Cell lysates and supernatants were collected 24 hours post treatment (hpt). To visualize ASC specks, LPS was diluted to a concentration of 5 µg/ml in Opti-MEM and cells were fixed 4 hpt. To activate the NLRP3 inflammasome BMMs were first primed with LPS in cell culture medium or left unstimulated for 3 h before cells were treated with 5 µM nigericin (Sigma-Aldrich) for 1 hour. At that time cell supernatants were collected for analysis by ELISA or LDH release assays. For monitoring autophagy, IFNγ primed and unprimed BMMs were washed with PBS, and then incubated for 2 hours in cell culture media, media containing Bafilomycin-A1 (100 nM, Sigma-Aldrich), or Hank’s balanced salt solution (HBSS, Thermo Fisher) prior to harvesting of cell lysates.

### Cytoplasmic delivery of LPS

Cells were primed overnight with 100 U/ml of IFNγ prior to LPS treatments. LPS was transfected into BMMs using Lipofectamine LTX with Plus Reagent (Thermo Fisher). BMMs were seeded in 96-well flat bottom plates at approximately 4 * 105 cells per well. For each well 375 ng of Lipofectamine LTX and 75 ng of LPS alongside 75 ng of Plus Reagent were each diluted in 2 µL of Opti-MEM. Following a 10-min incubation period at room temperature (RT), solutions were mixed and incubated for an additional 30 min at RT before the total reaction volume was brought up to 50 µL with media. Solutions were added to wells and plates centrifuged at 700 x g for 10 min. Cells were then incubated at 37°C for indicated times and cytoxicity assessed via LDH release assay. For co-delivery with *L. monocytogenes*, bacteria were grown overnight at 30°C, and then diluted to give an MOI of 10 in Opti-MEM. LPS was added at indicated concentrations and mixtures spun onto cells at 700 x g for 10 min. Cells were incubated at 37°C for 1 h, gentamicin was added to a final concentration of 15 µg/ml and cells were returned to the incubator for an additional 3 h before LDH release was assessed.

### LPS internalization and fluorescence associated cell sorting

Cells were treated with 2.5 µg/ml Alexa Fluor 488 conjugated LPS (L23351, Thermo Fisher) or unlabeled control LPS for indicated times. Following treatment, cells were washed twice with PBS, and collected in PBS containing 2% FBS. Fluorescence intensity data was collected a BD FACSCanto II system (BD Biosciences) and analysis was carried out using FlowJo Software.

### Cytotoxicity, viability and cytokine measurements

LDH release was measured using the CytoTox One homogenous membrane integrity assay (Promega). Cytotoxicity was calculated as a function of LDH release using the following formula: (sample – untreated control) / (100% lysis control – untreated control) * 100. In some instances, cell viability was determined using the CellTiter-Glo Luminescent Cell Viability Assay (Promega) and changes in viability relative untreated or uninfected cells was used to calculate cell death. Measurements of propidium iodide (PI) uptake were used to assess cell death over time. In these instances, treatments were performed in Opti-MEM containing 6 µg/ml PI and fluorescence measured at indicated time points. Relative fluorescence was calculated by subtracting PI fluorescence at time of initial treatment. For all cytotoxicity and viability assays, measurements were performed via Enspire 2300 (PerkinElmer) Multilabel Reader.

Cytokine concentrations were determined via enzyme-linked immunosorbent assay (ELISA) or Luminex Multiplex Assay as indicated in figure legends. ELISAs were performed per manufacturer’s instructions. The following ELISA kits were used: IL-1β Mouse ELISA kit (ThermoFisher), IL-18 Mouse ELISA kit (ThermoFisher), and ELISA MAX Deluxe Set Mouse TNFα (Biolegend). Measurements were performed via Enspire 2300 (PerkinElmer) Multilabel Reader. Luminex Multiplex Assay (Mouse 20-plex; Invitrogen) was performed in the Immunology Unit of the Regional Biocontainment Laboratory at Duke using a BioPlex 200 instrument.

### RNA isolation and quantitative PCR

RNA was isolated using a RNeasy Mini Kit (Qiagen) and reverse transcription accomplished using an iScript cDNA synthesis kit (Bio-Rad). Quantitative PCR was performed with QuantaFast SYBR Green PCR mix (Qiaqen) on a 7500 Fast Real-Time PCR system (Applied Biosystems). The following primers were used:

> mGAPDH F, 5′-GGTCCTCAGTGTAGCCCAAG-3′;
>
> mGAPDH R, 5′-AATGTGTCCGTCGTGGATCT-3′;
>
> mIL-1β
>
> F, 5’-GCAACTGTTCCTGAACTCAACT-3’;
>
> mIL-1β
>
> R, 5’-ATCTTTTGGGGTCCGTCAACT-3’;
>
> mTNF F, 5’-CATCTTCTCAAAATTCGAGTGACAA-3’;
>
> mTNF R, 5’-TGGGAGTAGACAAGGTACAACCC-3’;

### Immunocytochemistry

For visualizing ASC specks, cells were fixed for 5 min with ice-cold methanol. Coverslips were then washed 3 times with PBS, and cells were blocked with PBS containing 5% Bovine Serum Albumin (BSA) for 30 min at RT Cells were stained with rabbit anti-ASC antibody (AG-25B-0006; Adipogen; 1:1000) diluted in blocking buffer overnight at 4°C. Coverslips were then washed three times with PBS containing 5% BSA and 0.1% saponin and cells were next incubated with anti-rabbit Alexa Fluor 568-conjugated secondary antibody (Invitrogen; 1:1000) and Hoechst for 1 hour at RT followed by an additional three washes. For visualizing Gm130 and Irgm2, cells were fixed for 15 min at RT with 4% (wt/vol) paraformaldehyde. Coverslips were washed 3 times with PBS and then blocked with 5% BSA and 0.3 M glycine in PBS-saponin (0.1%) for 30 min at RT. Cells were stained with mouse anti-Gm130 (610823; BD Transduction Laboratories; 1:200) and rabbit anti-Irgm2 (1:500) diluted in blocking buffer for 1 hour at RT. Coverslips were then washed 3 times with PBS-saponin and stained with anti-mouse Alexa Fluor 488 and anti-rabbit Alexa Fluor 568-conjugated secondary antibodies (Invitrogen), and Hoechst for 1 hour at RT. Imaging was performed using a Zeiss Axioskop 2 upright epifluorescence microscope, a Zeiss 780 upright confocal microscope, or a Zeiss 880 Airyscan Fast Inverted Confocal. The custom-made polyclonal rabbit anti-Irgm2 antibody was raised against the N-terminal peptide KEAVESPEVKEFEYFS.

### Immunoblotting

Whole-cell lystates were harvested in RIPA lysis buffer (Sigma-Aldrich) containing 1 x protease inhibitor cocktail (Sigma-Aldrich). For visualizing cleaved caspase-1, supernatants were collected and protein concentrated via trichloroacetic acid (TCA) precipitation. Samples with mixed with a 100% (wt/vol) TCA solution in water to a final TCA concentration of 10% and incubated on ice overnight. Samples were then centrifuged at 21,130 x g for 10 min. Following centrifugation, supernatant was discarded, and pellets washed twice with ice-cold acetone. Pellets were air dried and resuspended in 8 M urea. Samples were loaded on 4-20% gradient SDS-PAGE gels and transferred to nitrocellulose or PVDF. Membranes were blocked for 1 hour at RT in Tris-buffered saline-0.1% Tween 20 (TBST) containing 5% BSA and incubated overnight at 4°C with indicated primary antibodies. Membranes were washed thrice with TBST and then incubated for 1 hour at RT with appropriate HRP-conjugated secondary antibodies. Membranes were washed and visualized on film. The following antibodies and dilutions were used: rabbit anti-Irm1 1B2 ((Butcher, Greene et al., 2005); 1:500), rabbit anti-Irgm2 (raised against peptide KEAVESPEVKEFEYFS; 1:500), rabbit anti-Irgm3 927B0 ((Taylor, Jeffers et al., 1996) 1:500), rabbit anti-GBP2 ((Coers et al., 2011);1:1000), rabbit anti-caspase-1 (AG-20B-0042; Adipogen; 1:1,000), mouse anti-β-actin (A2228; Sigma-Aldrich; 1:1,000), rabbit anti-P62 (PM045; MBL; 1:1,000), rabbit anti-LC3 (2775; Cell Signaling Technology; 1:500), goat anti-CD14 (AF982; Novus Biologicals; 1:500), and rat anti-Casp4 (17D9; Novus Biologicals; 1:500).

## Statistical analysis

Statistical analysis was performed using GraphPad Prism 7.00 software. Significance was determined using one-way analysis of variance (ANOVA), or two-way ANOVA with multiple comparisons tests as indicated in figure legends. Log rank Mantel-Cox test was used for determining significance in survival experiments. *p < 0.5, **p <0.01, ***p < 0.001. N.S, non significant.

## ACKNOWLEDGEMENTS

This work was supported by National Institutes of Health grants AI103197 (to J.C.), and AI148243 (to G.A.T and J.C.) We would like to thank Mr. She Zhe for his contributions to the production of the *Irgm2*-/- mouse as well as Ms. Cheryl Bock and Mr. Kucera at the Duke Cancer Institute Transgenic Mouse Facility (supported by NIH grant P30 CA014236) for their technical support in generating the *Irgm2*-/- as well as the pan*Irgm*-/- mouse lines. We are grateful to Dr. Masahiro Yamamoto for sharing the *GBP*chr3-/- mice with us. Luminex assay was performed in the Immunology Unit of the Regional Biocontainment Laboratory at Duke, which received partial support for construction from the National Institutes of Health, National Institute of Allergy and Infectious Diseases (UC6-AI058607). J.C. holds an Investigator in the Pathogenesis of Infectious Disease Award from the Burroughs Wellcome Fund, M.K. is supported by a German Research Foundation (DFG) fellowship, and R.F. was supported by a National Science Foundation (NSF) Graduate Research Fellowship.

## AUTHOR CONTRIBUTION

R.F. and J.C. conceived the project with input from G.A.T. R.F., J.D., M.K., G.W., A.S.P., A.F.H. and S.L. conducted the experiments and acquired data. R.F., J.D., M.K., G.W., S.L. and J.C. analyzed data. R.F. and J.C. wrote the manuscript. All authors contributed to manuscript reviewing and editing. N.O-R., J.M., M.J.K., S.W. and G.A.T. provided reagents. J.C. and G.A.T. supervised the project and acquired funding related to the project.

## EXPANDED VIEW (EV)

**Figure EV1.**
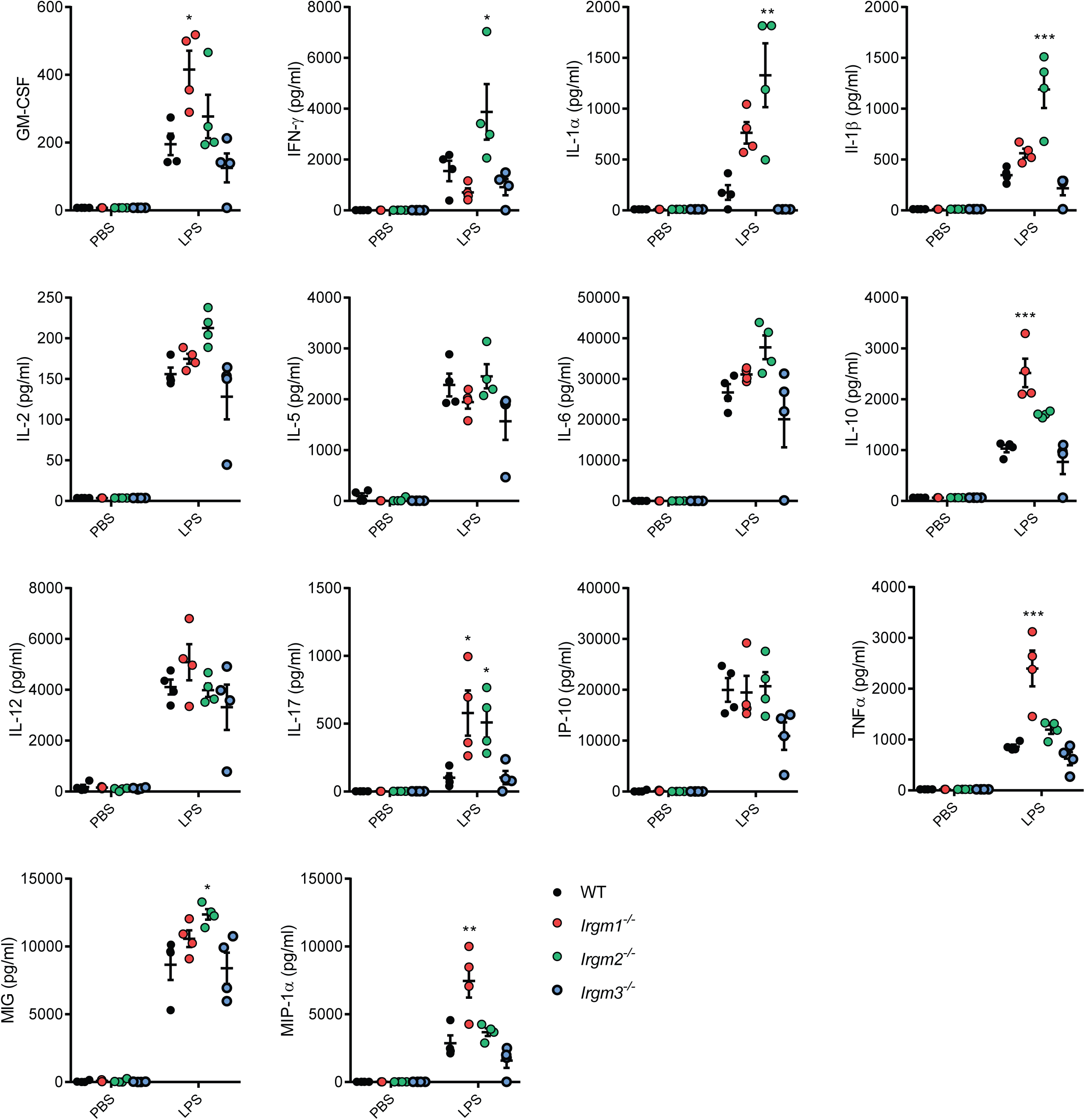
Absolute quantification of multiplex cytokine data of PBS- and LPS-treated WT and Irgm-deficient mouse strains. WT, *Irgm1*-/-, *Irgm2*-/- *and Irgm3*-/- mice were injected *i.p.* with LPS (8 mg/kg in PBS) or PBS alone. Serum was collected 4 hpi and concentration of indicated cytokines determined via Luminex platform (these data are shown normalized to WT mice in Fig 1A). n = 4 mice/genotype for all groups, except *Irgm1*-/- + PBS where n = 3 Data information: Data shown are means ± SEM. *p < 0.05, **p < 0.01, ***p < 0.001 for comparison between wildtype and indicated genotype by one-way ANOVA with Dunnett’s multiple comparison test.

**Figure EV2.**
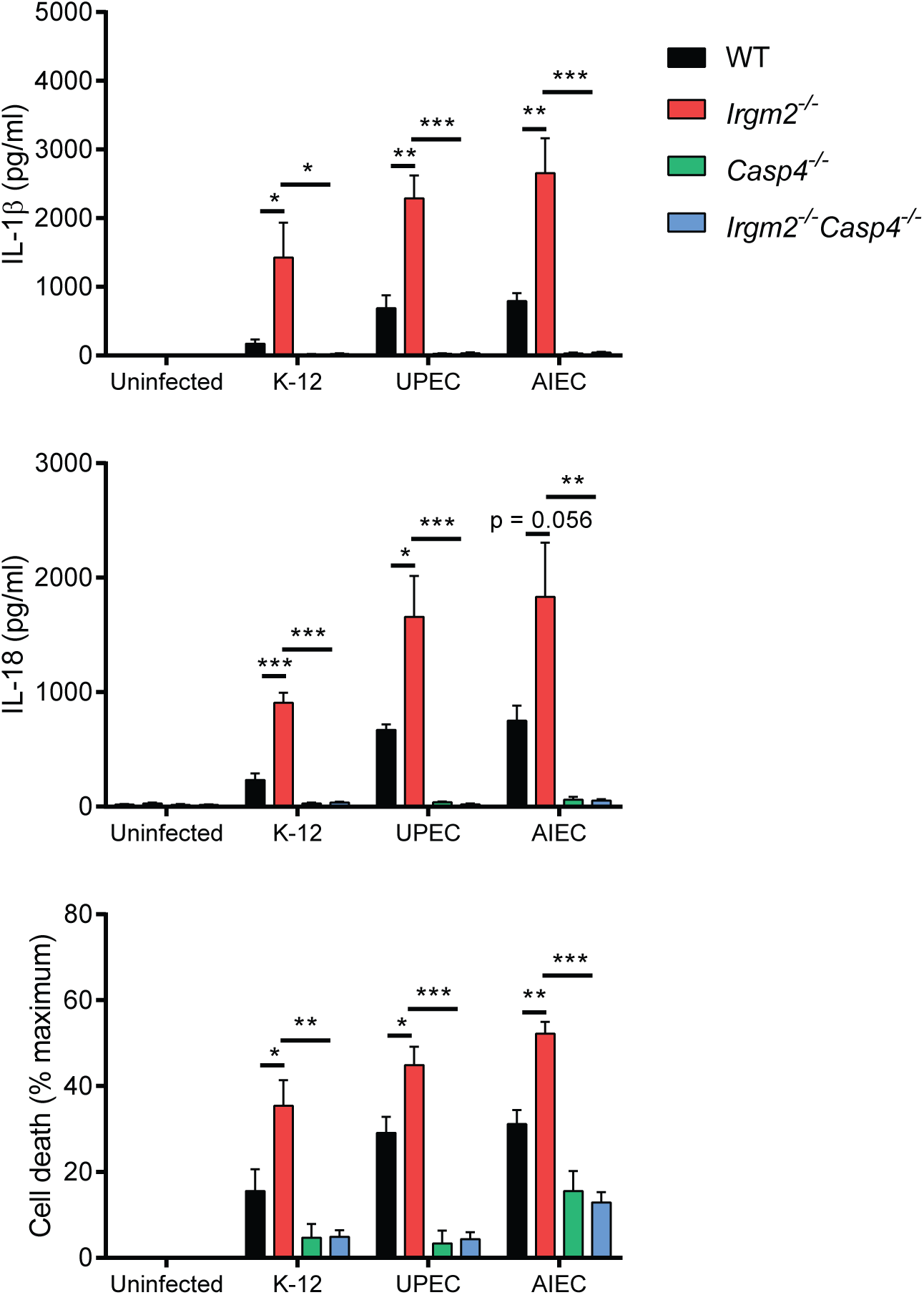
Irgm2 suppresses caspase-4 activation in response to pathogenic *E. coli* infections. WT, *Irgm2*-/-, *Casp4*-/- and *Irgm2*-/- *Casp4*-/- BMMs were infected with *E. coli* K-12, *coli* UPEC, or *E. coli* AIEC (MOI 25) and IL-1β, IL-18, and LDH release were assessed at 24 hpi. Shown are means ± SEM from n = 3 (IL-1β, IL-18) or n = 4 (LDH release) independent experiments. *p < 0.05, **p < 0.01, ***p < 0.001 for indicated comparisons by two-way ANOVA with Tukey’s multiple comparisons test.

**Figure EV3.**
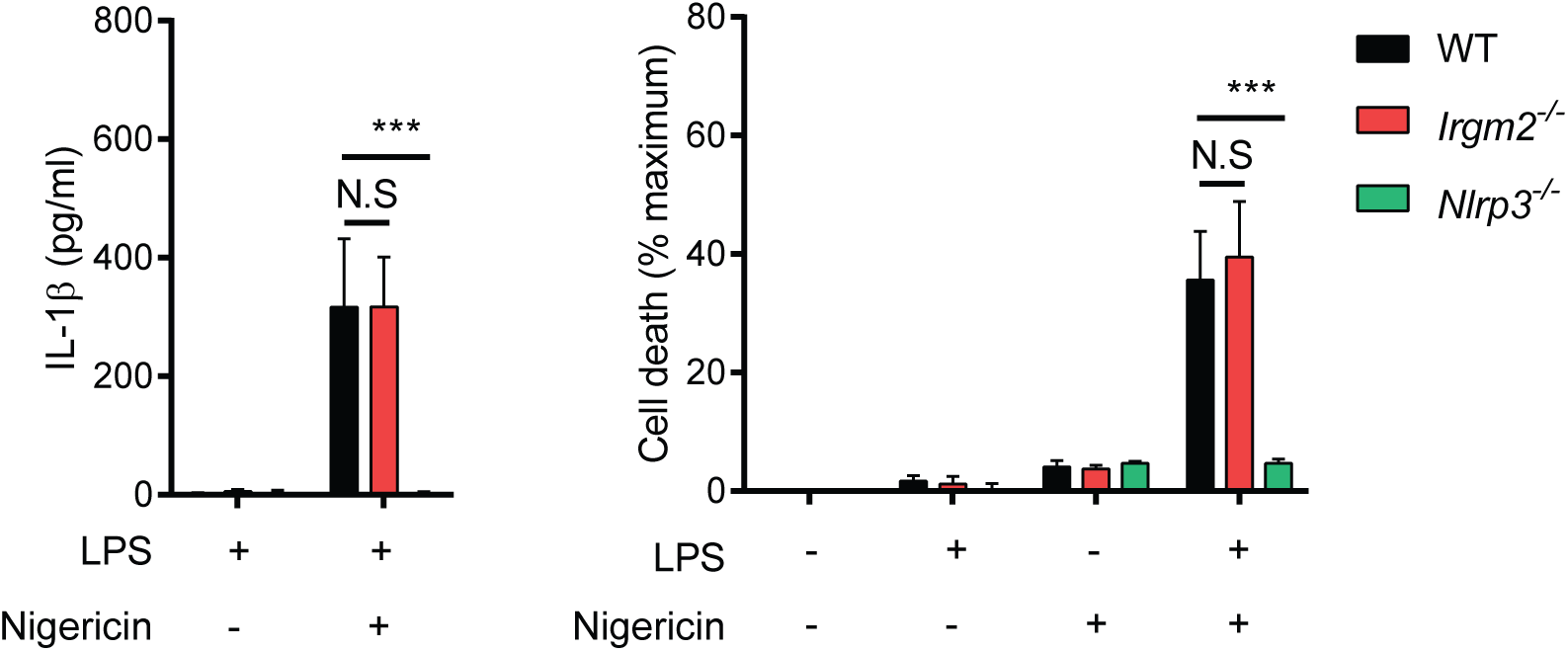
Nigericin-stimulated NLRP3 activation remains unchanged in Irgm2-deficient macrophages. WT, *Irgm2*-/-, and *Nlrp3*-/- BMMs were treated with LPS (0.1 µg/ml) for 3 h followed by nigericin for 1 h and IL-1β/LDH release was measured. Shown are means ± SEM from n = 5 independent experiments. ***p < 0.001 for indicated comparisons by two-way ANOVA with Tukey’s multiple comparisons test. N.S, non significant.

**Figure EV4.**
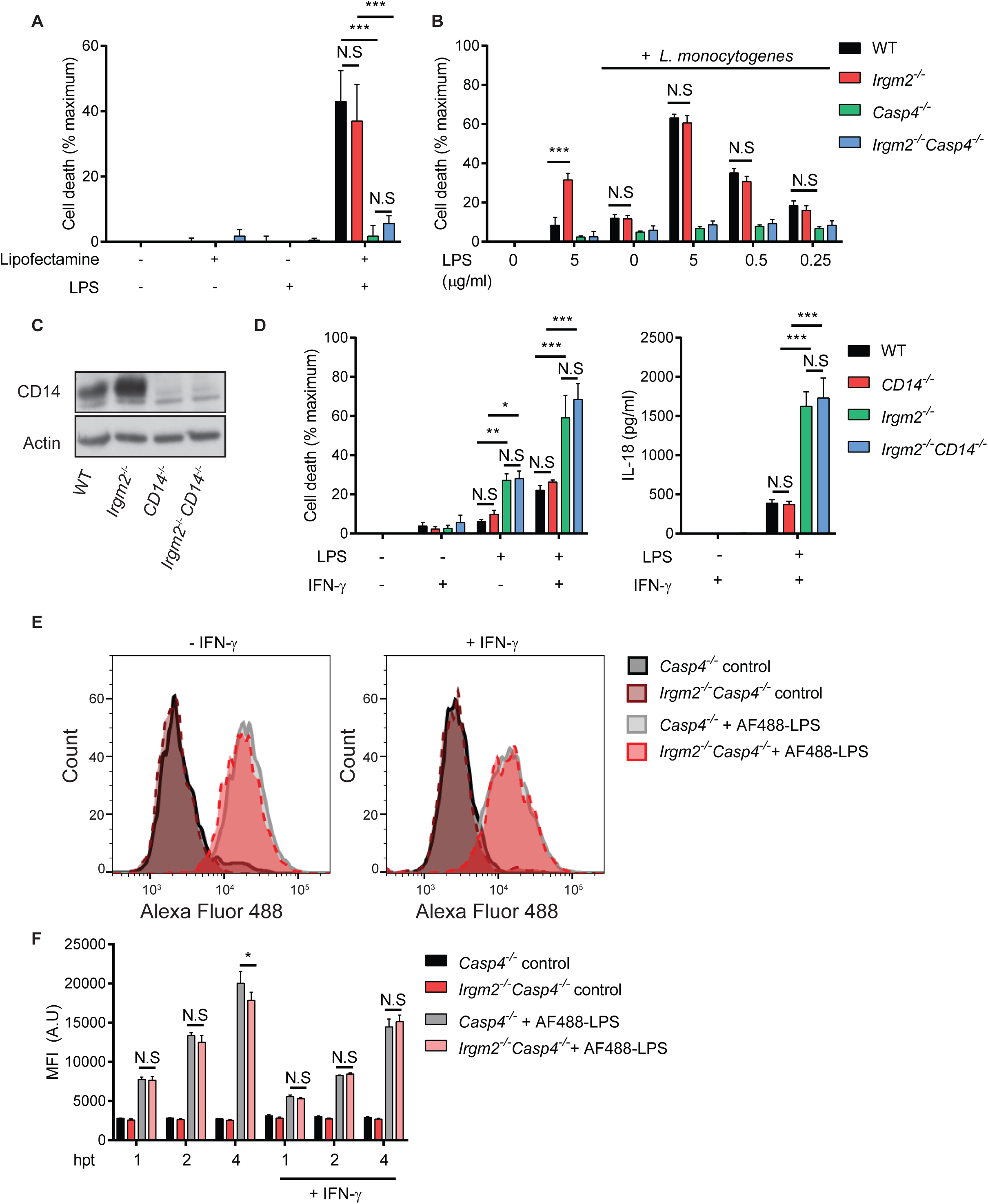
Irgm2 controls caspase-4 activation upstream from cytosolic LPS sensing. **(A)** IFNγ-primed WT, *Irgm2*-/-, *Casp4*-/- and *Irgm2*-/- *Casp4*-/- BMMs were transfected with LPS using lipofectamine LTX and LDH release was measured at 2 hpt. **(B)** IFNγ-primed WT, *Irgm2*-/-, *Casp4*-/- and *Irgm2*-/- *Casp4*-/- BMMs were co-treated with LPS (indicated doses) and *Listeria monocytogenes* (MOI 10) and LDH release measured at 4 hpt. **(C)** Lysates from WT, *CD14*-/-, *Irgm2*-/-, and *Irgm2*-/- *CD14*-/- BMMs were assessed for CD14 and actin protein levels via immunoblotting. **(D)** IFNγ-primed and unprimed WT, *CD14*-/-, *Irgm2*-/-, and *Irgm2*-/- *CD14*-/- BMMs were treated with LPS (1 µg/ml) for 24 h and LDH and IL-18 release measured. **(E)** IFNγ-primed and unprimed *Casp4-/-* and *Irgm2*-/-*Casp4-/-* BMMs were treated with Alexa Fluor 488-conjugated LPS or unconjugated LPS (control) for 4 hours and Alexa Fluor 488 cell fluorescence measured via FACS. Representative FACS data are depicted **(F)** IFNγ-primed and unprimed *Casp4-/-* and *Irgm2*-/-*Casp4-/-* BMMs were treated with Alexa Fluor 488 conjugated LPS or unconjugated LPS (control) for 1, 2, or 4 h and fluorescence measured via FACS. MFI (A.U) = Mean fluorescent intensity (arbitrary units). Data information: Data shown are means ± SEM from n = 3 (A, B, E, G) independent experiments. *p < 0.05, **p < 0.01, ***p < 0.001 for indicated comparisons by two-way ANOVA with Tukey’s multiple comparisons test. (D) is representative of two independent experiments. (E) are represents one of three independent experiments. N.S, non significant.

**Figure EV5.**
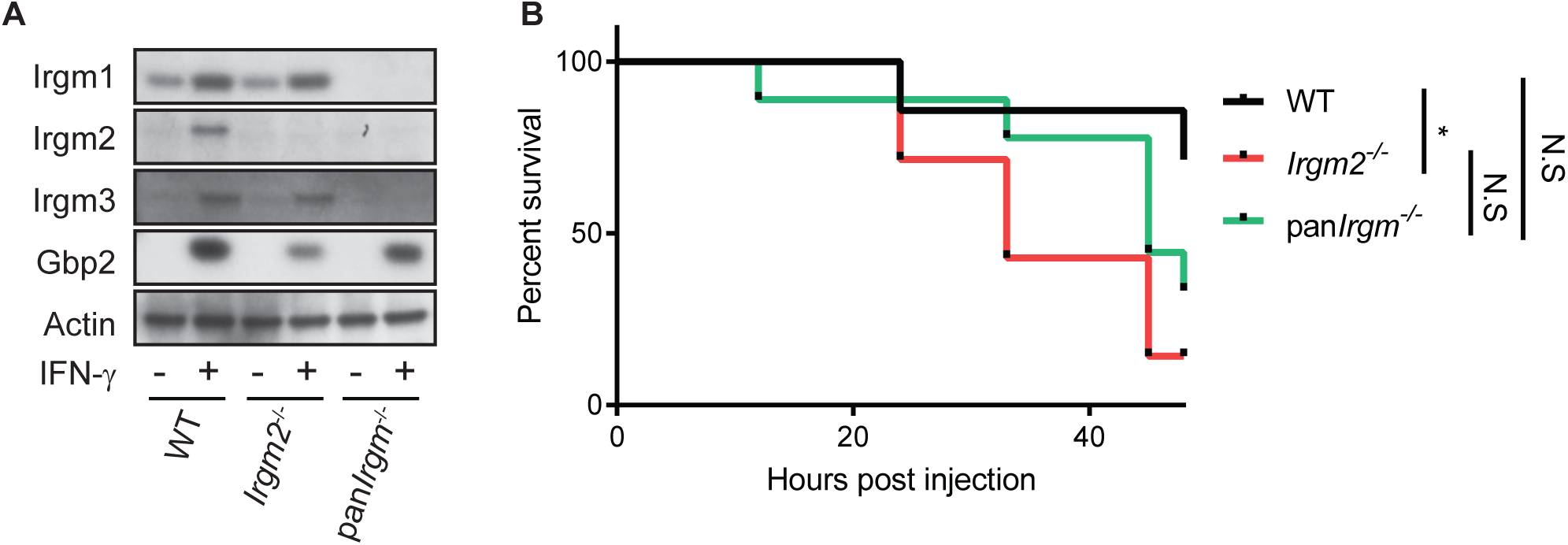
Pan*Irgm*-/- mice lacking expression of Irgm1/m2/m3 display improved viability during endotoxemia compared to *Irgm2*-/- mice. **(A)** WT, *Irgm2*-/-, and pan*Irgm*-/- BMMs were stimulated overnight with IFNγ or left untreated and cell lysates were collected. Lysates were assessed for Irgm1, Irgm2, Irgm3, Gbp2, and actin protein levels via immunoblotting. **(B)** WT (n = 7), *Irgm2*-/- (n = 7), and pan*Irgm*-/- (n = 9) mice were injected *i.p.* with LPS (2 mg/kg bodyweight). Morbidity and mortality were observed for 48 hours at 3 hour intervals. *p < 0.05, for indicated comparisons by log-rank Mantel-Cox test. N.S, non significant.

## SUPPLEMENTARY INFORMATION

**Figure S1.**
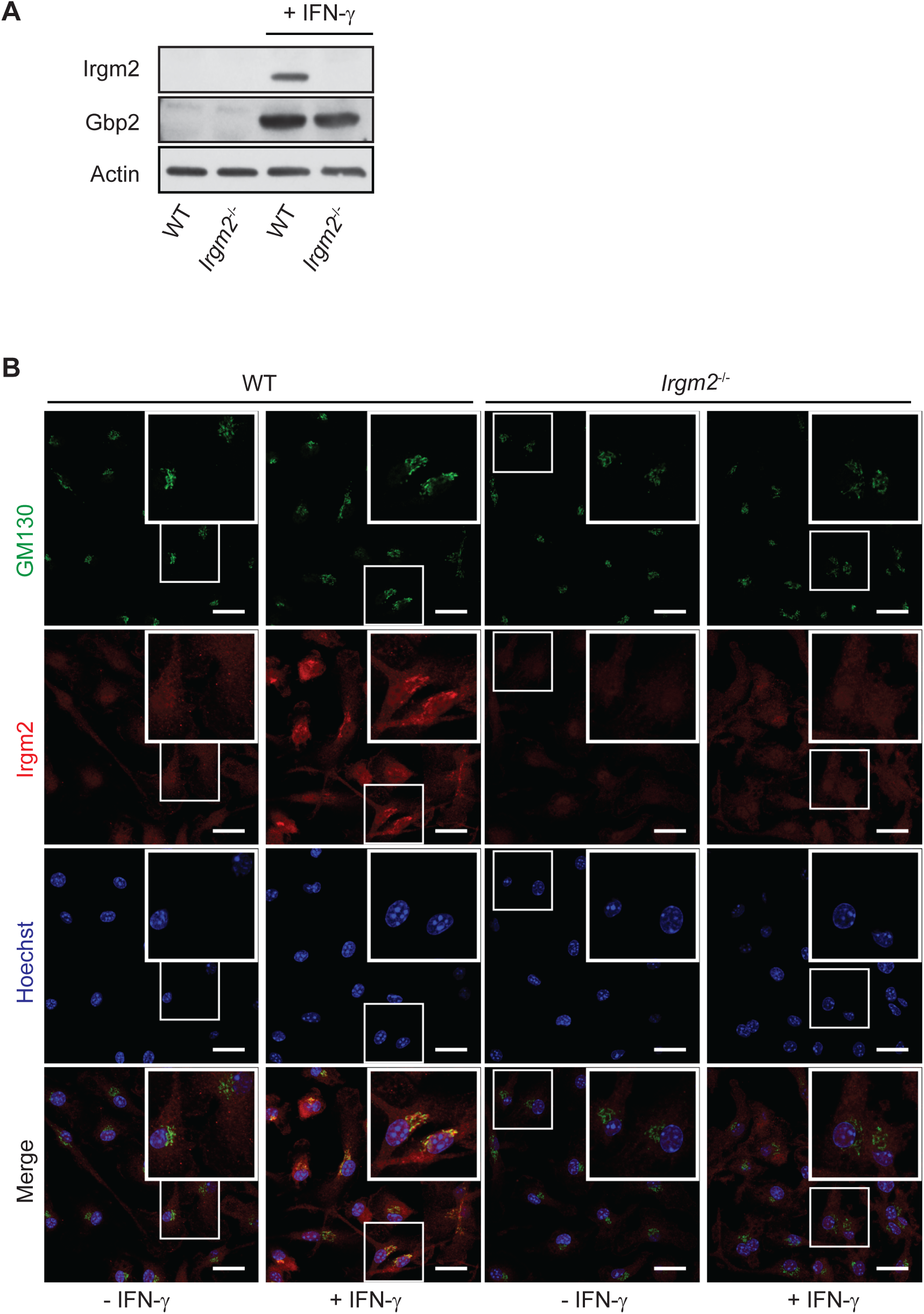
*Irgm2*-/- mice lack expression of the Golgi-resident Irgm2 protein. **(A)** WT, and *Irgm2*-/- BMMs were stimulated overnight with IFNγ or left untreated and cell lysates were collected. Lysates were assessed for Irgm2, Gbp2, and actin protein levels via immunoblotting. **(B)** IFNγ-primed and unprimed WT, and *Irgm2*-/- BMMs were fixed and subsequently stained with anti-GM130, and anti-Irgm2 antibodies as well as Hoescht. Representative images are shown.

**Figure S2.**
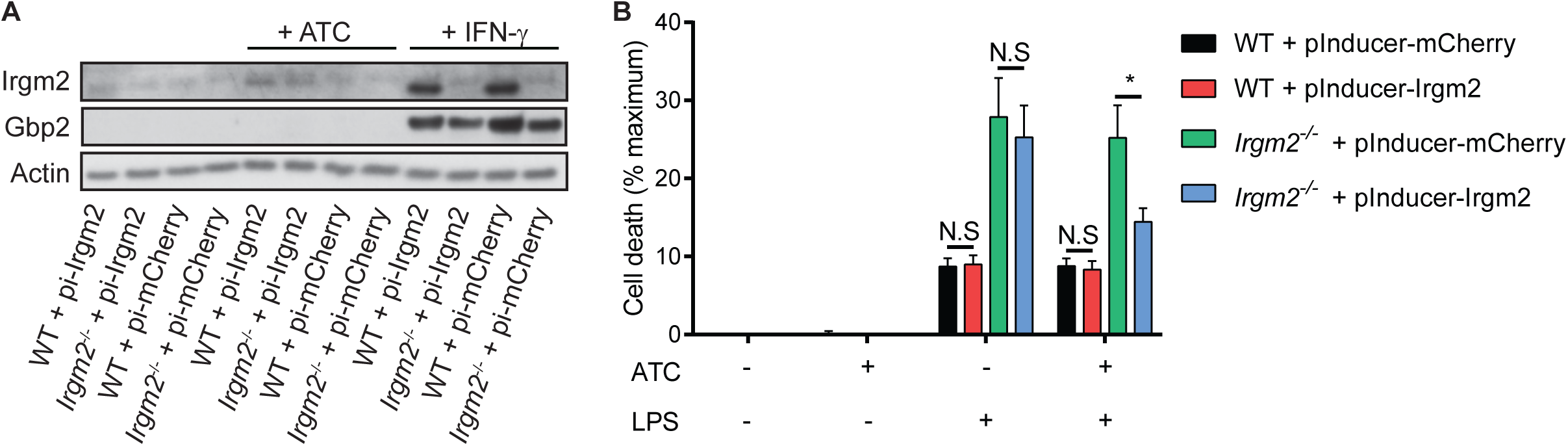
Ectopic expression of Irgm2 in *Irgm2*-/- iBMMs restores cell viability to WT levels. **(A)** WT and *Irgm2*-/- iBMMs stably expressing pInducer-mCherry or pInducer- Irgm2 were stimulated with aTc or IFNγ overnight. Cell lysates were assessed for Irgm2, Gbp2, and actin protein levels via immunoblotting. **(B)** WT and *Irgm2*-/- iBMMs stably expressing pInducer-mCherry or pInducer-Irgm2 were treated overnight with aTc and IFNγ or IFNγ alone followed by LPS (1 µg/ml) for 8 h at which time LDH release was measured. Means ± SEM are shown for n = 5 independent experiments. *p < 0.05 for indicated comparisons by two-way ANOVA with Tukey’s multiple comparison test. N.S, non significant.

**Figure S3.**
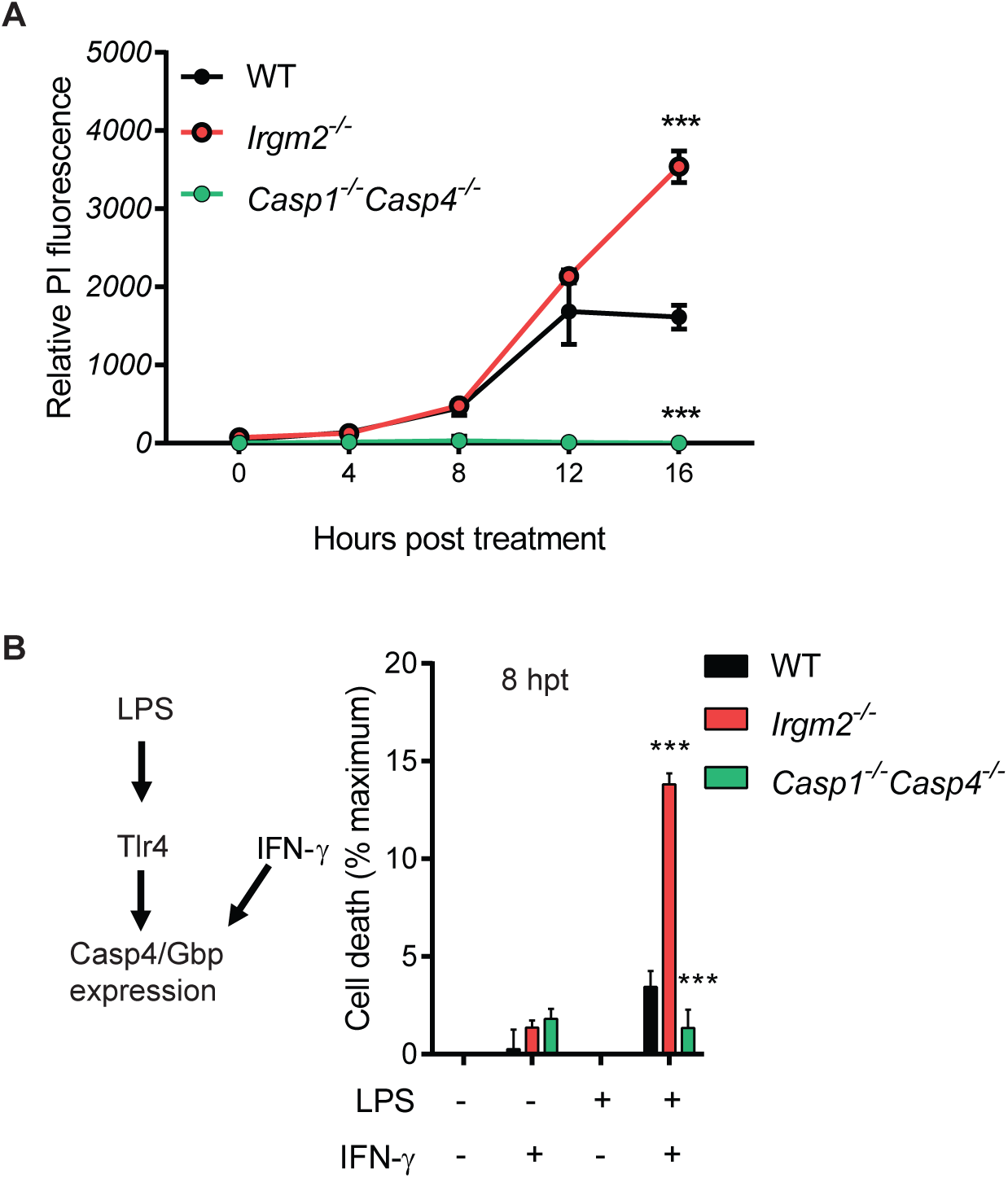
IFNγ priming accelerates the kinetics of pyroptotic cell death in *Irgm2*-/- BMMs. **(A)** WT, *Irgm2*-/- and *Casp1*-/-*Casp4*-/- BMMs were treated with LPS (1 µg/ml) and propidium iodide fluorescence assessed at indicated time points. **(B)** IFNγ-primed and unprimed WT, *Irgm2*-/- and *Casp1*-/-*Casp4*-/- BMMs were treated for 8 h with LPS (1 µg/ml) and LDH release was measured. Data information: Data shown is means ± SEM are shown for n = 3 independent experiments. *p < 0.05 for comparison between WT and indicated genotypes by two-way ANOVA with Tukey’s multiple comparison test. N.S, non significant.

